# The accelerating exposure of European protected areas to climate change

**DOI:** 10.1101/2024.11.25.625168

**Authors:** Marta Cimatti, Valerio Mezzanotte, Risto K. Heikkinen, Maria H. Hällfors, Dirk Nikolaus Karger, Moreno Di Marco

## Abstract

All ecosystems are affected by climate change, but differences in the pace of change will render some areas more exposed than others. Such spatial patterns of risk are important when assessing the continued functionality of protected area (PA) networks or planning for their expansion. Europe is undertaking an expansion of the PA network to cover 30% of its land and sea surface, but this must account for climate risk. Here, we estimate four metrics of future climate risk across Europe – local velocity, distance velocity, magnitude, and residence time – and assess the level of climate exposure of European PAs *vs* non-protected control sites. We also evaluate the intensity of climate risks on >1,000 European species of conservation concern, associated with Natura2000 sites. Our results show large spatial differences in climate change exposure across Europe, with faster pace and farther shifts in climate in the Boreal, Steppic, and Pannonian regions but slower changes in the Mediterranean, Alpine, Artic, and Macronesian regions. Climate change magnitude was higher for the Alpine, Mediterranean, and Steppic regions, implying large local differences between present and future climate. These spatial risk patterns were largely consistent across scenarios, but with up to three times higher risk under the most pessimistic *vs* the most optimistic scenario. Large variation in climate exposure for species of conservation concern was revealed, including 11 species which are highly dependent on Natura2000 sites and predicted to experience rapid climate change. Our results provide guidance for managing European PAs, and expanding their coverage, by pinpointing areas offering more stable climates. We emphasize the need for connectivity across the network, to support species adaption via range shifting. This is especially the case in areas facing high climate change magnitude but low climate velocity, implying that climate conditions similar to current ones will be found nearby.

## 2. Introduction

Biodiversity and ecosystems support human societies worldwide, but the exploitation of natural resources and conversion of natural land for human use is causing widespread biodiversity loss (Díaz et al., 2019; Dirzo et al., 2014). On top of these direct pressures, climate change is projected to cause accelerating impacts on biodiversity (Dawson, 2011; Díaz et al., 2019; Urban, 2015; Wiens & Zelinka, 2024), intensifying the impacts of other pressures such as land-use change (Brook et al., 2008; Heikkinen et al., 2021; Mantyka-Pringle et al., 2012; Mantyka-Pringle et al., 2015; Oliver et al., 2015). The primary means to halt the loss of biodiversity has been the establishment of protected areas (PAs), and other area-based conservation interventions (Margules & Pressey, 2000; Watson et al., 2014). PAs counteract or slow down (Santangeli et al., 2023) the overall loss of biodiversity and the fragmentation of ecosystems, and form the backbone of many conservation strategies worldwide (S. Hoffmann et al., 2018; Pimm et al., 2014). Protection of areas also represent the only conservation mechanism to preserve the last remaining habitat of many rare and threatened species (Lawrence et al., 2021; Nila & Hossain, 2019; Pacifici et al., 2020). Thus, the Kunming-Montreal Global Biodiversity Framework calls for an expansion of “*protected areas and other effective area-based conservation measures - covering at least 30% of the planet by 2030*”. In line with this framework, the EU Biodiversity Strategy for 2030 commits member states to protect at least 30 % of land and sea by 2030, with 10 % of this area placed under strict protection.

In 2024, European protected areas covered 26.1% of the land surface, with the Natura 2000 network, the largest PA network in the world (EEA, 2024; Eurostat, 2022) covering 18.6% of the area. Natura 2000 offers substantial protection for species of conservation concern, including (but not limited to) those listed in the Birds Directive and Habitats Directive (Trochet & Schmeller, 2013), although gaps in its coverage remain. In fact, a large number of threatened species of mammals, birds, butterflies, and reptiles are substantially dependent on PAs, and especially Natura 2000 sites for survival (Sluis et al., 2016; but see Ricci et al. (2024). However, European PAs, like most PA networks around the globe, have been established without proper consideration of the risks stemming from climate change (S. Hoffmann, 2021; Lai et al., 2022). Climate change threatens the effectiveness of PAs in preserving biodiversity, as species might be unable to cope with new climatic conditions and need to shift their ranges to track their preferred climate, or alter their phenology, reproductive behaviour, or ecology (Araújo et al., 2011; Bellard et al., 2012; S. Hoffmann et al., 2019; Mendez Angarita et al., 2023).

As a response to climate change, terrestrial species are moving poleward and/or towards higher elevations (Bellard et al., 2012; Hällfors et al., 2024; Scheffers et al., 2016; Thomas & Gillingham, 2015), while aquatic species are also moving deeper (Brito-Morales et al., 2020). Thus, climate change threatens to compromise the effectiveness of PAs in protecting biodiversity, as species might be driven out of PAs (Lehikoinen et al., 2021; Loarie et al., 2009; Santangeli et al., 2017). To accommodate the needs of biodiversity under a changing climate, it is crucial to assess the projected climatic exposure of PAs and associated risks to species persistence (Lai et al., 2022).

In response to rapidly accelerating climate change, there is accelerating interest in assessing climate risks using metrics such as velocity and magnitude (Brito-Morales et al., 2018; Garcia et al., 2014). Several global studies have analysed the rate of predicted climate change in PAs (Elsen et al., 2020; S. Hoffmann et al., 2019; Loarie et al., 2009). A number of studies have also focussed on climate risk in European terrestrial habitats and PAs (Araújo et al., 2011; Brito-Morales et al., 2018; Heikkinen et al., 2020; Lai et al., 2022; Nila & Hossain, 2019). Yet, despite the increased use of climate change metrics in basic and applied PA studies, we still have an uncomprehensive understanding of how climatic risk that can be measured through a variety of metrics (and in addition multiplied by several potential scenarios, and several climate models) will develop, and how these affect PAs *vs* control sites of similar characteristics. Developing on this front would allow an improved understanding of the potential implications for species of conservation concern.

Here we evaluate climate risk for Europe and its PAs at a 1km-resolution, using four metrics of climate change, three future emission scenarios, and three earth system models. We evaluate whether some regions are more highly exposed than others according to each metric, whether climate risks are higher inside *vs* outside PAs (using a statistical matching technique), and finally, assess the level of risks for species of conservation concern. The four metrics of climate change used here represent complementary aspects of risk (Brito-Morales et al., 2018; Loarie et al., 2009; Williams et al., 2007): (1) local-based velocity, describing the intensity of change in a given location relative to local variation in climatic conditions; (2) distance-based velocity, describing the distance between current climate in a location and the closest location with analogue conditions in the future; (3) magnitude of change, depicting how much the climatic conditions of a location change over time; (4) climate residence time, which indicates how long current climatic conditions will persist within the boundaries of an area (in our case, a PA).

These four metrics provide different and complementary information on how far and how rapidly climate analogues will move and how much current conditions will change in specific locations or within PAs. It is essential to use them in combination as different regions, PAs, and species occurrence sites are differentially exposed to the climate change and the risk that is measured by each metric describe different aspects of change (Garcia et al., 2014; Ordonez & Williams, 2013).This assessment is thus a crucial step in climate-proofing conservation management and planning, helping conservation planners to ensure climatic resilience of new PAs and corridors within the expanded Trans-European Nature Network (TEN-N).

## 3. Materials and methods

We examined the spatial variation of four complementary metrics of climatic exposure across Europe: local (gradient based) and distance (analogue based) climate velocity, magnitude of climate change, and residence time. For each metric, we compiled climate change data under nine combinations of three climate scenarios (SSP1-2.6, SSP3-7.0, SSP5-8.5) and three earth system models (ESMs): GFDL-ESM4, UKESM1-0-LL, MPI-ESM1-2-HR. This entailed a total of 36 climate risk predictions (4 metrics * 3 scenarios * 3 ESMs; Table S1). For each climate metric and climate scenario, we also generated an ensemble prediction across ESMs, by averaging data from the three models, and retrieved the standard deviation.

Our study area covers the territories of EU 46 extended from Greenland in the west to the Urals in the east, including Turkey and parts of North African countries to cover the entire mediterranean shoreline (Fig. 1-S1-S3, S6-S7). However, the study area for the propensity scores matching includes countries from EU 28 and Belarus, Bosnia and Herzegovina, Moldova, Montenegro, North Macedonia, Norway, Serbia, Ukraine, and the United Kingdom (Fig. S4).

**Figure 1.**
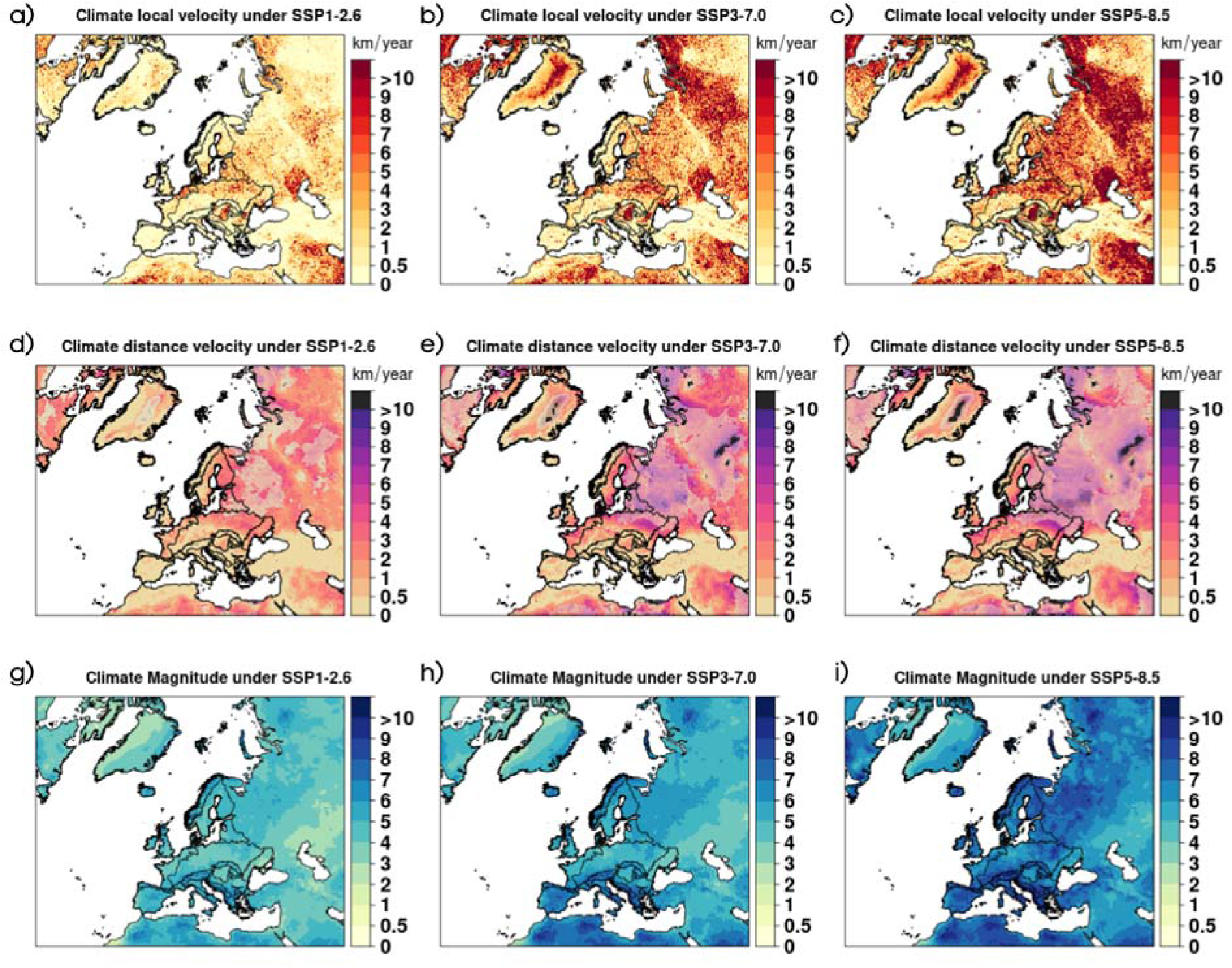
Estimates of different climatic metric values at 1 km resolution under scenarios SSP1-2.6, panes to the left (a), d), and g)), SSP3-7.0 in the middle (b), e), and h)), and SSP5-8.5 on the right (c), f), and i)) based on temperature and precipitation (km/yr). Panels a), b), and c) on the upper row show values for the absolute sum of local velocity. Panels d), e) and f) in the second row show the distance (analog based) velocity, where areas in shaded colours depict raster cells for which there was no climate analogue within the 500km buffer around the focal cell for one (lighter shade), two (darker shade) or all three ESMs (in black, see Fig.2). Panels g), h), and i) on the bottom row show values for climate magnitude (intensity of the change, adimensional). Metric values for individual climatic variables and ESM are shown in Figs. S6-7.

### 3.1 Selection of climate variables, scenarios, and models

We calculated climate change metrics based on two bioclimatic variables from CHELSA V2.1 (Karger et al., 2017, 2021) describing general climatic conditions: mean annual near-surface (2m) air-temperature (*temperature*) and the annual precipitation rate (*precipitation*). We chose this reduced set of climate variables to represent overall climatic risk, in line with previous studies (Asamoah et al., 2021; Loarie et al., 2009).

Using an ensemble of ESMs for different climate change scenarios is commonly recommended for mastering the range of possible future climates and associated uncertainties (Pereira et al., 2010; Thuiller et al., 2019) stemming from different equilibrium climate sensitivity of ESMs (Knutti et al., 2017). We selected a set of scenarios from the Coupled Model Intercomparison Project Phase 6 (CMIP6) that encompass low to high levels of future carbon emission (O’Neill et al., 2016). In scenario SSP1-2.6 “Sustainability – Taking the green road” an increasing shift toward sustainable practices is envisioned with persisting efforts to limit global warming to below 2°C compared to pre-industrial level. Scenario SSP3-7.0 “Regional Rivalry – A Rocky Road” involves strong land use change and entails the highest methane and air pollution precursor emissions with global mean temperature increases of 1.95 - 4.38°C by 2100 (Tebaldi et al., 2021). Scenario SSP5-8.5 “Fossil-fuelled Development – Taking the Highway’’ represents a world with a strong push for economic and social development at the expense of climate mitigation, with a temperature increase of 2.40 - 5.57°C by 2100 (Tebaldi et al., 2021). In parallel, we use three earth system models (ESMs) following the priority proposed by the Inter-Sectoral Impact Model Intercomparison Project Phase 3 (ISIMIP) (Frieler et al., 2024; Lange, 2019) to represent inherent uncertainty around climatic modelling approaches. This priority accounts for process representation, structural independence, equilibrium climate sensitivity, and model performance in the historical period.

### 3.2 Selection of climatic metrics

Based on each combination of climate scenarios and ESMs resulting, we calculated four climatic metrics to quantify the exposure of each PA to climate change.

The first metric is local velocity (*sensu* Loarie et al., 2009) which is calculated based on local climatic gradients (Carroll et al., 2015; Loarie et al., 2009). The local velocity of climate change represents the rate of movement of climatic isopleths across a landscape (Loarie et al., 2009), describing the speed and direction at which a given organism needs to move to stay under similar climatic conditions. From an ecological perspective, local velocity provides estimates of species adaptation potential by finding suitable conditions adjacent to their current occurrence sites in subsequent years. We measured the velocity of change following the gradient-based approach (Loarie et al., 2009), and using an automated workflow based on the R functions ‘tempTrend’ and ‘spatGrad’ in the ‘gVoCC’ package (García Molinos et al., 2019). The velocity of change is defined as the ratio between the temporal trend of a climate variable (the rate of change of the variable through time, for baseline (1980-201) and future (2041 – 2070) periods (estimated as a regression slope), and the corresponding local spatial gradient of that variable (i.e. the vector sum of longitudinal and latitudinal pairwise differences at each focal cell using a 3 x 3-cell neighbourhood), described in Eq. [1].

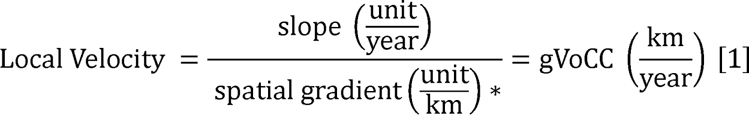

\**unit/year*: (°C/km for temperature and mm/km for precipitation)

The second metric was the distance climate velocity, which is based on geographical distances between locations of current climate classes and their future analogues (Brito-Morales et al., 2018; Hamann et al., 2015). From an ecological perspective, this metric provides a measure of how far species need to move within a given time frame to find similar conditions to those currently experienced. Distance (analogue-based) climate velocity is calculated as the distance, *d,* to the geographically closest climate analogue for each focal cell divided by the time elapsed, *t,* between baseline (1981 −2010) and future periods (2041 – 2070), calculated according to Eq. [2].

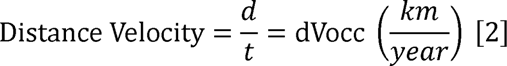

Climate analogues were identified using multivariate (minimum) Euclidean distance between current and future climate, using a dissimilarity threshold of 1.5 standard deviations based on the variability of historical climate conditions (Doxa et al., 2022; García Molinos et al., 2017). Calculating velocity through analogues can pose computational challenges since it involves scouring the entire extent of the study area to find the spatially closest cell with the same climatic attributes of each focal cell (i.e. solving an N^2^ problem, considering N= 29632989 cells). Thus, we limited the search radius around each focal cell to 500 km, as a compromise between computational time and biological significance.

The third metric, climate change magnitude, measures change in a climate parameter over time at a given location, providing information on the extent of the change in each location in absolute terms of the climate variable (Garcia et al., 2014). Ecologically, this metric provides direct information on the climate change to which a species is exposed in a given location (Williams et al., 2007; Williams & Jackson, 2007). We quantified magnitude as dissimilarities between baseline (1981 −2010) and future (2041 – 2070) climate by using the standardized Euclidean distance (SED) (Gavin et al., 2003; Williams et al., 2007), as in Eq. [3].

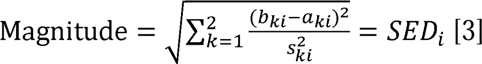

In Eq. [3] *a_ki_* and *b_ki_* are the baseline and future means for climate variable *k* at locations *i*, respectively, while *s^2^_ki_* is the standard deviation of the interannual variability for 1981 – 2010. The SED equally weights all variables and highlights predictions for 2041 – 2070 that are large relative to inter-annual variability during 1981 – 2010.

As our fourth metric, we assessed the *climate residence time* of European PAs. Residence time is a metric that describes after how long (e.g., in years) a specific isoline will cease to exist in a focal area (Brito-Morales et al., 2018; Loarie et al., 2009). Ecologically, a short residence time means a given PA might soon become unable to sustain suitable condition for the biodiversity it currently hosts (Heikkinen et al., 2020). In our analysis, we measured residence time as the time after which future climatic conditions disappear from within the PA. We calculated the residence time based on *temperature* and *precipitation* using the VoCC package and the resTime function (García Molinos et al., 2019), according to Eq. [4]:

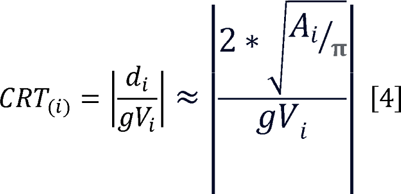

where *di* is the diameter of a PA’s polygon *i*, which is calculated based on the areal size of the polygon (*Ai*), and *gVi* is the average climate velocity measured within the PA polygon *i*.

### 3.3 Comparing climatic risk inside *vs* outside PAs

We created a spatial map of European PAs by extracting data from the World Database on Protected Areas (WDPA, IUCN & UNEP-WCMC, 2023). For every PA, key information is provided, such as spatial data, its extent, the year of establishment, and the IUCN category of protection level. The database contains multiple duplicates of the same areas (Vimal et al., 2021), one for each legislative instrument protecting them. To remove such duplicates and eliminate overlapping polygons we used the *wdpa_clean* function in the R package *wdpar* (Hanson, 2022) and assigned each PA to the highest IUCN protection category among those covered by several legislative instruments. We also generated circular buffers with areas matching the reported size around protected areas that were listed as points but lacked polygonal boundaries. We included in our analysis only terrestrial sites, clipping PAs by the coast using the country borders’ polygons from the Global Administrative Areas Dataset (GADM, https://gadm.org/).

To compare climate change inside *vs* outside PAs, we identified non-protected “control” sites using a propensity score matching approach (Geldmann et al., 2019; Negret et al., 2020; Stuart et al., 2013). Propensity score matching is defined as the probability of receiving treatment, given specific covariates, which allows unbiased matching between subjects in the control condition and subjects who have “undergone treatment” (Olmos & Govindasamy, 2015; Stuart et al., 2013). For every protected focal cell (i.e. our “subjects”), the propensity score matching selects a non-protected cell with the most similar baseline environmental characteristics to provide an unbiased correspondence for comparison. We defined a cell as “protected” (i.e., a subject receiving the treatment) if it overlapped by >50% with a PA, and a cell as “non protected” (i.e., a control subject) if it overlapped by <5% with a PA. Cells with marginal protection (between 5 and 50%) were excluded. We used several environmental variables in the matching process (Table S2): the 19 bioclimatic variables from CHELSA V.2.1 (Karger et al., 2017, 2021), land cover from Copernicus (Buchhorn et al., 2020), Human Footprint (Venter et al., 2016), and topography derived from the SRTM elevation data (Farr et al., 2007). Before performing the matching, all variables were standardized, using the scale function in R (R Core Team, 2023). Analyses were run at a 1 km^2^ resolution reprojecting all the variables in ETRS89 Lambert Azimuthal Equal Area (ETRS_LAEA).

Matching was performed individually for each European biogeographical region (only including states with exclusively European territories) using the “MatchIt” package in R (Ho et al., 2011), with the nearest neighbour method without replacement. A threshold of SD<0.25 was used to exclude low-quality matches, where the distance in propensity scores between treatment and control was too low (Cuenca et al., 2016; Negret et al., 2020).In the last step, we used a Wilcoxon test to assess whether significant differences existed between the climate change metrics of protected and unprotected areas in each biogeographic region (exact matching).

In order to assess the quality of matching, we checked for covariate balance before and after the matching within the original dataset and in the matched sample. To assess the balance between the treated and control groups, we used the standardized mean difference test (SMD) and the C-statistic test. These tests allow comparing the distribution of baseline covariates between treatment groups in observational studies and as a balance measure of individual covariates before and after matching (Rahman & Islam, 2021). Here, we used the “Cobalt” package (Greifer, 2024), and set a cut-off at a standardized difference of 10%. SMDs close to zero indicate a good balance (Austin & Austin, 2009; Haukoos & Lewis, 2015; Rahman & Islam, 2021), with recommended values of SMDs being below 0.1. The C-statistic ranges from 0.5 to 1.0, with the minimum indicating that the propensity score model is not able to discriminate between treated and untreated units after matching, indicating a perfect balance of covariates (Franklin et al., 2014).

### 3.4 Climatic exposure for species within Natura 2000 areas

We estimated climate change exposure for species occurring in the Natura 2000 sites to assess the potential biodiversity implications within this PA network. We produced a list of species that meet two criteria: they are (I) identified as vulnerable to climate change according to the Threat Classification Scheme of the IUCN Red List (IUCN, 2023) and (II) they are reported to occur within one or more Natura 2000 sites. In this case, using Natura 2000 sites, instead of all PAs, allowed us to select all species with verified presence in each site (information which is not systematically available for other PAs). This resulted in 1,011 species encompassing vertebrates, invertebrates, and plants. We measured the mean and median values for each climate metric for all Natura 2000 sites in which at least one such species occurred.

For 514 species with spatial distribution information available from the IUCN Red List, we also measured the percentage overlap of the distribution with the Natura 2000 network. This allowed us to calculate the climatic metrics weighted according to the size of the PAs. Our aim was not to measure climate “risk” separately for each species, which would often require consideration of other factors such as species’ sensitivity and adaptability (Foden et al., 2013). Instead, we use this to demonstrate the level of climate exposure that already vulnerable species are projected to face within protected sites.

## 4. Results

Overall, we found notable spatial differences in climate change exposure across Europe for all four climatic metrics considered, as well as among the three climate scenarios studied. We found that, under scenario SSP1-2.6, the European median local (gradient based) climate velocity would decrease from a present (1981 − 2010) value of 1.77 km/year across the study area to a future value of 0.73km/year (sd = 0.62; Fig. S5a-S5d). The situation was inverse under high-emission scenarios, with a median future velocity of 3.22 km/year under SSP5 (s.d. = 1.15 km/year). Distance climate velocity (i.e. distance to analogue climate between 1981 − 2010 and 2041 – 2070) increased in high-emission scenarios compared scenario SSP1-2.6. However, for the distance velocity it was not always possible to find an analogue climate within the 500 km buffer around the focal cell. In particular, under the most extreme ESM (UKESM1-0-LL) and the most pessimistic emission scenarios we found that large portions of the study area (up to 50% of the area under SSP5) had no future analogue within 500km (especially in the Northeastern parts; Fig S2d-S2e-S2f). For this reason, we do not report median distance velocity values, as this would likely underestimate climate risk for large parts of our study area.

The median climate magnitude (i.e. dissimilarity between 1981 − 2010 and 2041 – 2070 climates) showed similar trends as velocity metrics, ranging from 3.90 for scenario SSP1-2.6 (s.d. = 2.49; Fig. S5c-S5-S5f) to 6.13 (s.d. = 2.61) under scenario SSP5–8.5 (Fig. 1).

As regards local velocity, the most exposed biogeographic regions were the Pannonian, Boreal, and Steppic, while for distance velocity the most exposed regions were the Steppic, Boreal, and Atlantic. For both velocity measures, the less exposed biogeographic regions were the Alpine, Mediterranean and Macaronesian. In contrast, the Alpine and Mediterranean regions experienced the highest values for climate magnitude, followed by the Artic, whereas the lowest magnitude values occurred in the Pannonian (Table S3).

### 4.1 Climate risk in PAs *vs* control sites

Our statistical matching process was successful in determining control sites for European PAs: the covariate imbalance was reduced from a mean C-statistic value of 0.744 before the matching to a mean value of 0.535 across all biogeographic regions after matching (Table S4). The standardized mean difference test also performed well for the whole of Europe (Fig. S8) since all covariates were adjusted through the matching and reached a good balance, within the 0.1 threshold. On a European level, the propensity score matching suggests that PAs are more exposed to climate change compared to unprotected sites, with higher median local and distance velocity, especially under the SSP5 scenario (Figs. 2 and S9-S10-S11, Tables S4-S5). Conversely, the climate magnitude measure showed an opposite pattern compared to climate velocities (Fig. 2, Table S4-S5), with control areas having higher values of magnitude than PAs. The Wilcoxon test confirmed that the values of climatic metrics between PAs and control areas significantly differ on the European level (Table S5).

**Figure 2.**
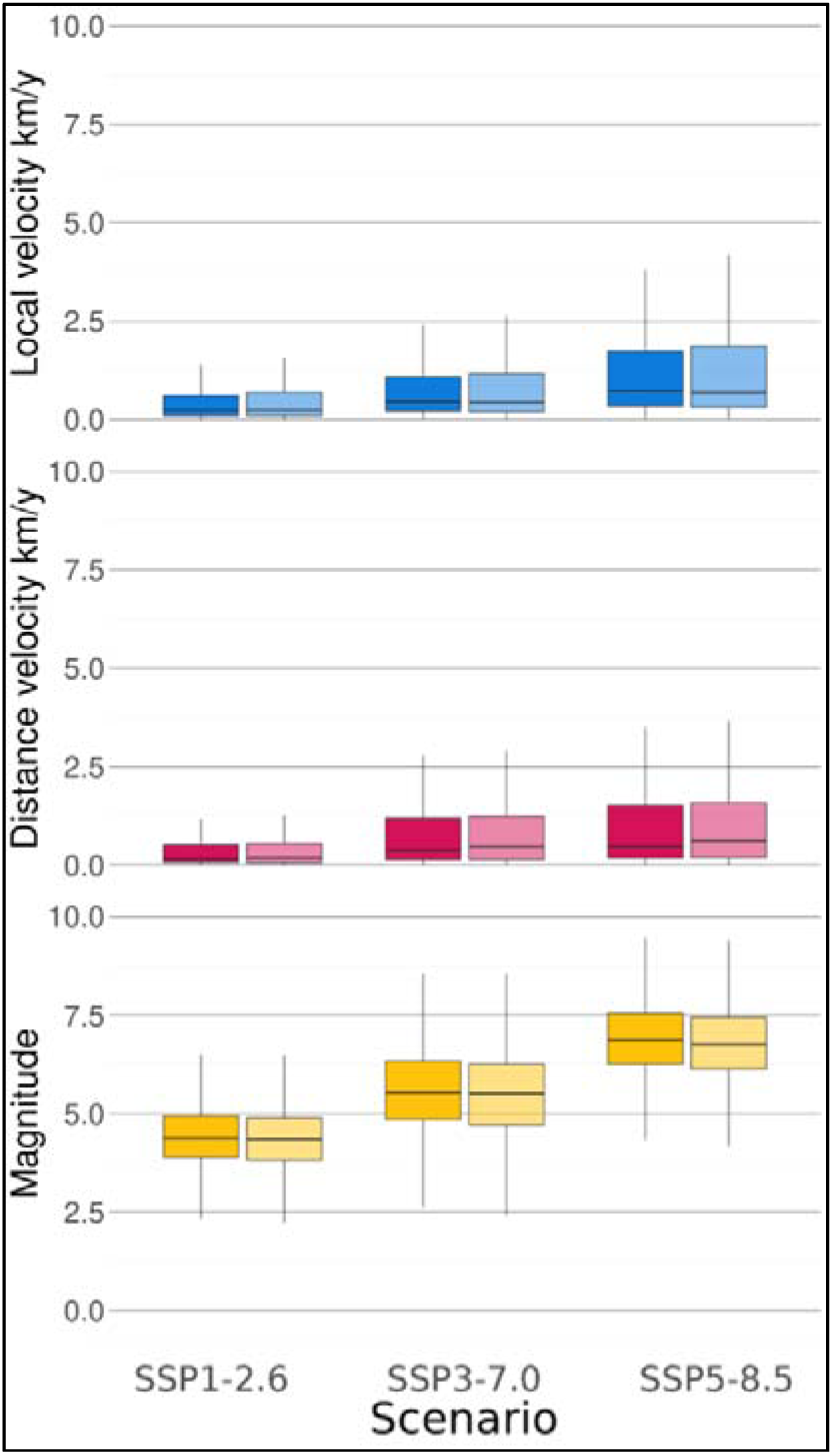
Boxplots showing the range of the three VoCC metrics: (I) local velocity (blue, absolute sum of the velocities for temperature and precipitation), (II) distance velocity (red), and (III) climate magnitude (yellow) inside PAs (lighter colour) and in control areas (darker colour) under different future climate scenarios. Results for distance velocity exclude grid cells from matching study areas (Fig. S4) for which it was not possible to find future analogue climates (16.5%) within the 3 SSP-RCP scenarios.

The trend for the climatic metrics in the 10 biogeographic regions generally followed the overall European-wide trend across the scenarios, with greater median values of local and distance velocity within PAs compared to control areas, and with climate magnitude showing a very small difference between PAs and controls (either higher or lower, depending on the region). In general, the differences in VoCC metrics between PAs and controls were statistically significant at the scale of biogeographic regions. However, there were some exceptions, e.g., for temperature velocity in the Black Sea region under all scenarios, and for climate local velocity in the mediterranean region under scenario SSP1-2.6 (Table S6). Moreover, PAs in different biogeographic regions showed different levels of climate change exposure (Fig. S9).

Under all scenarios, the highest values of local climate velocity occurred in the Pannonian, Steppic, and Boreal biogeographic regions (Fig. 1, Fig. S9), while Alpine, Mediterranean, and Macaronesian biogeographic regions showed the lowest values. The median value for local velocity of temperature (Fig. S10) increased under each scenario compared to the baseline period, with the highest values found under the SSP5-8.5 scenario. Nevertheless, there was a high degree of spatial heterogeneity (Fig. S10), and the highest values were found in the Pannonian and Steppic regions. For precipitation (Fig. S11), local velocity was sometimes negative, indicating decreasing precipitation which can lead to increasingly dry conditions in association with increased temperature. The Steppic, Pannonian, and Black Sea region were estimated to be most affected by drier conditions.

As previously mentioned, result for distance velocity exclude cells for which it was not possible to find cells with analogue climate (50% of the study area, but just 16.5% of EU 28 territory included in the propensity score matching).In the EU 28 area, the Boreal, Steppic, and Continental regions were overall the most exposed regions based on distance velocity values, while the Mediterranean and Alpine regions were less exposed. The differences in distance velocity values between PAs and control regions follow the overall European trend, with higher distance velocity values in PAs compared to control regions, except for the Steppic and Pannonian regions.

The general trend for climate magnitude was a dramatic increase within PAs across all biogeographic regions, when going from the lowest emission scenario to the highest. The Alpine region showed the highest value of climate magnitude across all scenarios, followed by the Mediterranean and the Arctic. In general, control regions showed higher values of magnitude compared to PAs, apart from the Artic, Boreal, Pannonian, and Steppic regions. While the magnitude of temperature follows the same pattern, the highest magnitude values for the precipitation were found for the PAs in the Boreal region (Figs. S9-10).

Overall, the differences in magnitude values between PAs and controls were all statistically significant. However, there were no significant difference for climate magnitude in the Black Sea region under any scenario, and none in the metrics based on precipitation in the Macaronesia region under any scenario (Table S6).

### 4.2 Residence time

The mean residence time across all PAs was 25.5 years, 12.2 years, and 8.0 years for scenarios SSP1–2.6, SSP3-7.0, and SSP5–8.5, respectively. The biogeographic regions with the highest residence time were the Arctic and the Alpine biogeographic regions, followed by the Mediterranean and Macaronesian regions (Figs. S12-S13, Table S7). In contrast, the Boreal, Pannonian, and Steppic regions showed the lowest residence times across all scenarios, with low residence times even in the low-emission scenario (median values < 5 years for SSP 1-2.6).

### 4.3 Climatic exposure for species in the Natura 2000 network

The 1,011 species identified as threatened by climate change and that occur within the Natura 2000 network showed similar climate change exposure values across the taxonomic groups (Figs. 3 and S18, Table S8). For each species, we estimated the median local climate velocity within its Natura 2000 sites of occurrence. We found that median local velocity of included species’ occurrences within Natura 2000 tripled (from 1.09 km/y to 3.45 km/y) when moving from scenario SSP1-2.6 to SSP5-8.5. Similarly, the magnitude increased by ca. 50% (from 4.25 to 6.75).

**Figure 3.**
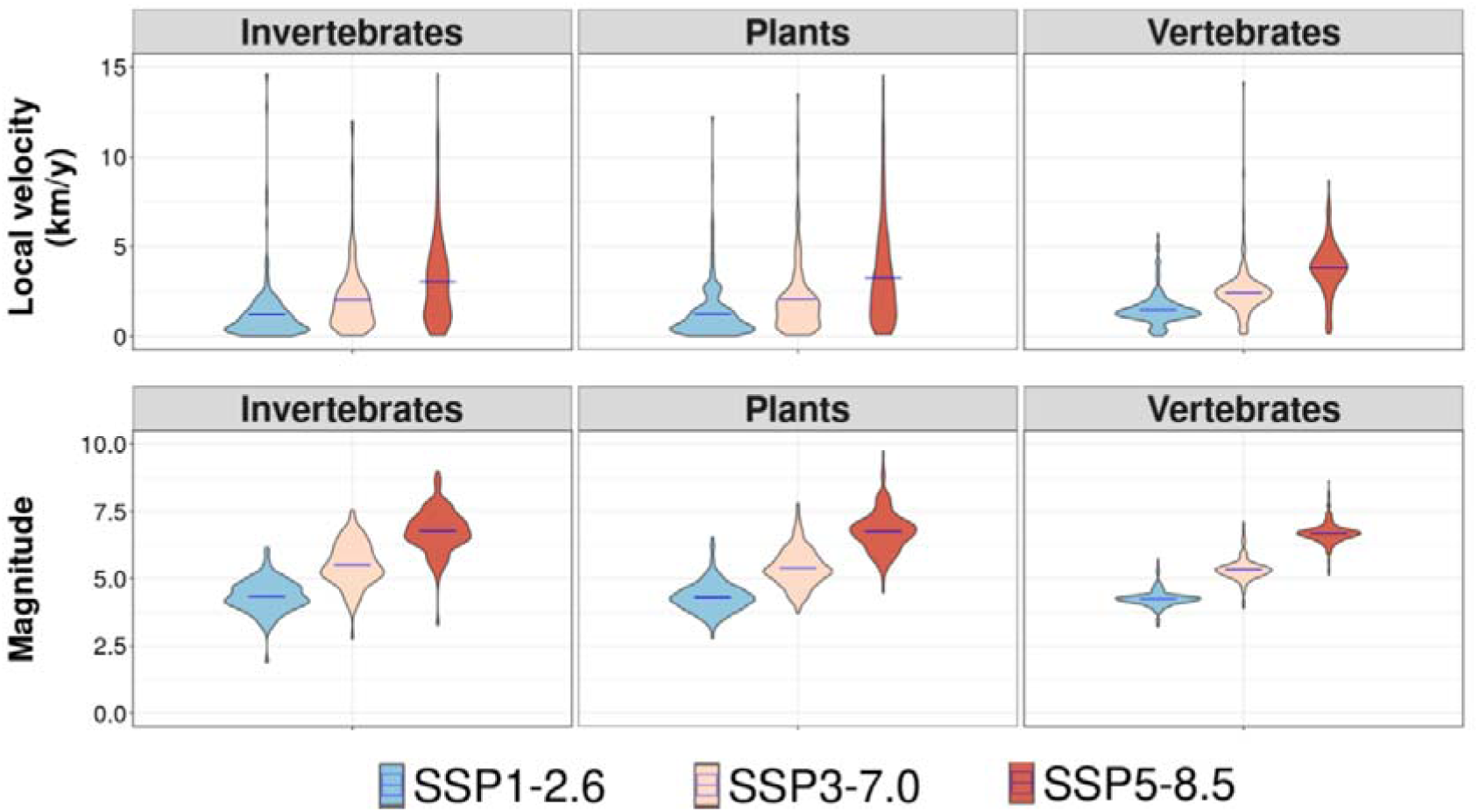
Predicted local velocity and magnitude for the 1011 threatened species occurring within the Natura 2000 network, presented by main taxonomic group.

For the 514 species with detailed spatial distribution information available from the IUCN Red List database, climate exposure within the Natura 2000 network was assessed in relation to overlap of the species’ distribution area with Natura 2000 sites (Figs. 4 and S19, Table S9). Climate exposure and distribution area overlap were poorly related, indicating that species with any level of representation within the Natura 2000 network are equally highly exposed to climate change risks (Figs. 4 and S19). All species groups showed a decreased future velocity under SSP1 compared to the historical period, but much higher velocities under scenarios SSP3 and SSP5 (Fig. 4). Eleven tetrapod species had >50% of their range within Natura 2000 sites, suggesting a high dependence on the Natura 2000 network (Table S10).

**Figure 4.**
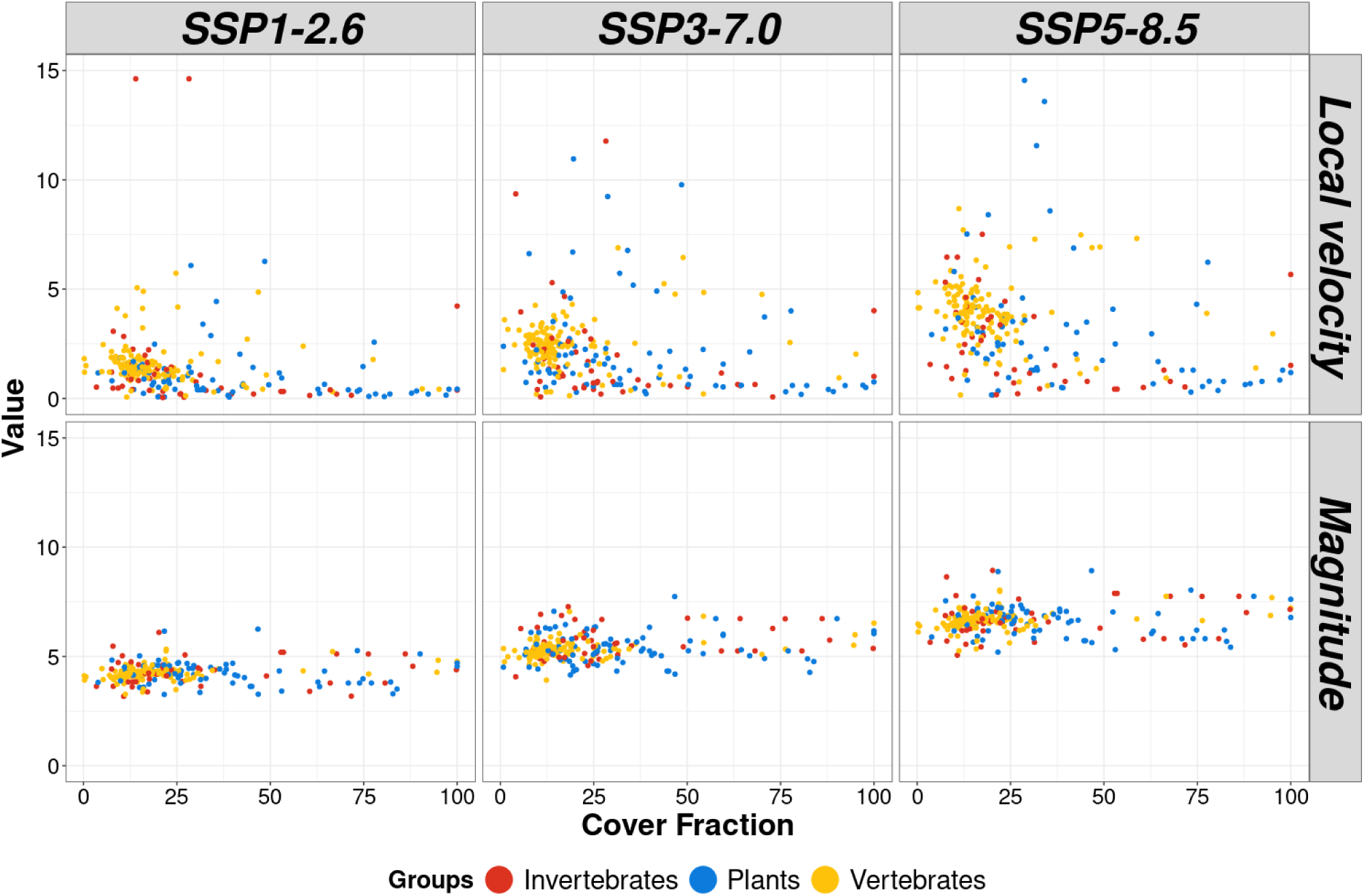
Predicted climate local velocity under SSP1-2.6 and SSP 5-8.5 scenarios compared to the cover fraction (overlap between species distributions and Natura 2000 sites) of invertebrate, plant and vertebrate (514 species)

## 5. Discussion

### 5.1 Geographic patterns across multiple dimensions of climate change

Biodiversity is becoming increasingly exposed to the impacts of climate change both inside PAs and in the intervening matrix, as shown previously at global (S. Hoffmann et al., 2019; Loarie et al., 2009), continental (Batllori et al., 2017; Belote et al., 2018) and regional scales (Barber et al., 2016; Hamann et al., 2015; Heikkinen et al., 2020; Stagl et al., 2015). Estimating climate velocity has become a standard approach for examining the exposure of biodiversity to climate change (Brito-Morales et al., 2018). However, to develop more comprehensive assessments of risks, multiple dimensions of climate exposure should be examined to obtain a more holistic picture that can also guide in PA expansion (Garcia et al., 2014; Nadeau & Fuller, 2015).

In Europe, the Natura 2000 network is projected to face major changes in climatic conditions, challenging its long-term performance in safeguarding valuable species populations and habitats (Lai et al., 2022; Nila & Hossain, 2019; Stagl et al., 2015). For example, Nila and Hossain (2019) showed that novel climate conditions may emerge in the future in Natura 2000 sites of the Alpine region. Also, several PAs in the Continental and Mediterranean regions are projected to experience new climates while current climatic conditions disappear (Nila & Hossain, 2019). In three iconic protected areas in Central Europe, Bierbza National Park, Balaton National Park, and Riesenferner-Ahrn Nature Park, monthly temperatures has been projected to increase by 3-3.7°C by 2071 – 2100, revealing systematic signals of warming across more than 20 investigated climate scenarios (Stagl et al., 2015). In Northern Europe, contrasting spatial velocity patterns have been projected, with some areas and PAs experiencing high exposure of winter *vs* summer-time temperature velocity, including a full-scale turnover of current temperature conditions in the PAs (Heikkinen et al., 2020). Lai et al. (2022) assessed the climatic exposure of the Natura 2000 network, defining velocity hotspots as areas facing both substantial forward (current to future) and backward (future to current) velocities (≥ 5km/year), and coldspots as areas with low forward and backward velocities (< 0.5km/year). Both high forward and backward velocities occurred particularly widely under a high emission scenario in the flatter parts of the Boreal and Continental regions and in high-elevation areas in the Alpine region. Coldspots were found in coastal Mediterranean areas and in low-elevation areas in the Alpine region. The integrated use of forward and backward velocities by Lai et al. (2022) provided complementary measures, occasionally showing contradicting metrics which indicates that the same species can be exposed to different climate change impacts in climatically marginal areas.

Our study goes beyond previous ones, by providing a many-sided high resolution (1km) assessment of climate exposure based on the latest climate data (those from CMIP 6) and employing a combination of complementary metrics, climate models, and scenarios. Our results for local velocities and distance velocities are largely congruent with the forward velocity results of Lai et al. (2022), suggesting that the highest exposure is found in inland Europe within both the Continental and Boreal regions. However, with our approach we were additionally able to recognise that the largest climate magnitude changes will occur in the Mediterranean region and in some parts of the Alpine region, but high climate velocities will only sporadically occur in the Mediterranean region. This is probably due to the high degree of spatial heterogeneity that is present inside this biogeographic region. Higher mean climate residence times within the Natura 2000 were projected to occur in the lowland areas of the Pannonian and Steppic regions and extensively across the Boreal region, showing both overlapping highly exposed areas but also some spatial differences, e.g., in the western part of boreal Fennoscandia.

Taken together, our findings highlight that there is substantial geographic variation in the level of climate exposure across areas in Europe, but also large variation depending on the metric considered. This may generate dissimilar pressures depending on species’ distributions across Europe. Corresponding differences have been revealed for contiguous USA where the highest forward and backward velocities overlapped only partly with local climate dissimilarity (magnitude of multivariate climate change; Belote et al., 2018), again highlighting the different dimension of climate change exposure underlying different metrics. This, together with our results, motivates the combined use of multiple metrics to reach a holistic understanding of climate change exposure.

Although the overall results show large variation across the study area and among the different metrics, we also identified certain areas where different dimensions of high climate risks coincide. Accounting for such joint climate risks is important for conservation planning as they can help identify elevated exposure to species residing in these areas (Belote et al., 2018; Lai et al., 2022). We found areas in North Africa, North East Europe, the eastern parts of the Continental and Boreal regions, and the Baltic stand out as high-exposure areas in terms of both velocity and magnitude of change under all scenarios. Most areas in the Mediterranean basin and Norway are hotspots of climate magnitude under both optimistic and pessimistic scenarios. In contrast, hotspots of local and distance velocity can be found in central Europe (Figs. S14-S15). We also identified areas where several metrics converged on a likely low risk of climate change exposure. These were especially prevalent in Cyprus, Lebanon, the Syrian coast, and large areas between the Caspian sea and Persian Gulf (Figs. S16-S17).

When PAs were compared to environmentally similar corresponding areas across Europe, our results revealed statistically significant differences between the mean values based on three climate metrics. On average, PAs showed lower climate change magnitudes, but higher local and distance velocities compared to unprotected areas. Similar results were obtained by Lai et al. (2022) for the Natura 2000 network based on forward velocity values. However, in our results, these differences were small, although statistically significant due to our very large sample size (1km grid cells). Nevertheless, if our results are assumed to reflect current and future risk, these modest Europe-wide differences may be of limited ecological relevance, suggesting that PAs and matrix areas are, overall, facing similar climate change threats. However, for climate-wise conservation planning, our velocity assessments provides crucial information, particularly by allowing the identification of areas with the highest or lowest velocities into which new conservation activities might be targeted (Hlásny et al., 2021; Lai et al., 2022).

For our fourth metric, residence time, there are notable (up to 20-fold) differences between PAs located in different biogeographic regions, with the Boreal, Pannonian, and Steppic regions showing the lowest and Alpine and Arctic regions the highest residence times (Figs S12-S13, Table S7). This means that in PAs situated in more extreme environments, current climatic conditions will prevail for a longer time than elsewhere. This is in line with earlier studies (see Brito-Morales et al., 2018 and citations therein) showing a clear linkage between low topographic heterogeneity and shorter residence times, suggesting elevated risks for species in such areas (Heikkinen et al., 2020; Huey et al., 2012; Loarie et al., 2009).

### 5.2 Species exposure to different dimensions of climate change

Developing multifaceted assessments of climate change velocity and magnitude is critical for understanding how climate change affects the exposure of biodiversity. Such assessments can guide adaptation needs and improve our understanding of the survival prospects of species, thereby facilitating climate-wise conservation planning. Both climate velocity and the magnitude of local changes contribute to reshuffling species distributions, creating novel climates, and altering ecological interactions (Brito-Morales et al., 2018; Gaüzère et al., 2023; S. Hoffmann et al., 2019; Lurgi et al., 2012; Williams & Jackson, 2007). However, to improve relevance, climate metrics need to be explicitly related to different species.

We found substantial variation in climate exposure among Natura 2000 species included in our analysis, regardless of their taxonomic group. The main differences were found among scenarios, with exposure values up to three times higher based on SSP 5-8.5 compared to SSP 1-2.6 (Fig. S18-S19). This result, however, must be interpreted with caution as different species have different levels of sensitivity and adaptive capacity to climatic change, regardless of their exposure to it (Foden et al., 2013). For example, diet breath and habitat specialisation determine range shift potential in birds and butterflies, while limited dispersal ability is known to affect amphibians and reptiles (Campbell et al., 2024; Lurgi et al., 2012; McMenamin et al., 2008). While climate velocity metrics offer a holistic understanding of exposure, it is also important to recognise that they encase a simplification that species are locally adapted to current climatic conditions in small spatial locations. In reality, many species have relatively broad climatic tolerances and can thrive in a wider set of climatic conditions than those prevailing in a limited number of spatially defined grid cells. Nevertheless, the four metrics used in our study represent different aspects of climate change exposure, each with unique implications for biodiversity conservation (Garcia et al., 2014; Ordonez et al., 2016), facilitating a diverse threat assessment for the species. Areas with low climate velocity, such as mid-elevation slopes in mountainous regions likely provide more stable settings. If climate analogues are found in adjacent areas, this can support species in their adaptive responses to climate change (Ackerly et al., 2010; Garcia et al., 2014). Local changes in climate conditions – here, climate magnitude exposure – can directly impact species’ physiology, morphology, and behaviour leading to demographic changes (Camacho et al., 2024; Khaliq et al., 2014). In particular, species populations that currently occur in conditions close to their climatic tolerance limits and species with specialized climatic requirements or limited adaptive capacity are at higher risk of declines or local extinctions under high exposure to climate magnitude (Araújo et al., 2013; Doucette et al., 2023; A. A. Hoffmann et al., 2013). Our results show the highest levels of climate magnitude in the Alpine and Mediterranean regions. In the latter region, changes in temperature together with even modest changes in precipitation can significantly jeopardise populations of species that are less tolerant to extended periods of drought and increased frequency of fires (Batllori et al., 2019). In the uppermost parts of the Alpine region, species are at risk of facing cul-de-sacs due to disappearing local climate space (Hamann et al., 2015). This, in combination with future climate analogues located far away, may be particularly detrimental to sessile and slowly dispersing non-volant species (Dobrowski & Parks, 2016; Lai et al., 2022).

In many parts of Europe, projected velocities for temperature exceeded 5km/year and sporadically even 10km/year. This pace can be considered critically high as it exceeds the rate of observed geographic shifts even for some highly mobile species, such as boreal birds (Heikkinen et al., 2021). For some less mobile species, e.g., dispersal-limited plant species with narrow climatic niches, high velocities clearly produce substantial obstacles for successful range shifts (Barber et al., 2016). Moreover, even species with higher dispersal capacities, ranging from mammals to bryophytes, have been predicted to lag behind future changes in climate (Lenoir et al., 2020; Schloss et al., 2012; Zanatta et al., 2020). The risks brought by climate change are elevated in the regions where PAs and suitable habitat patches occur as isolated fragments in human-dominated and ecologically degraded landscapes (Asamoah et al., 2021). In such areas, higher velocities may increase species range fragmentation and complicate their ability to follow suitable climates (Brito-Morales et al., 2018; Garcia et al., 2014).

The climate residence times define for how long stable and suitable climate conditions remain in the PAs for species that are dependent on habitats found therein. Residence times were found to be smallest in the lowlands of Continental and Boreal Regions, with median values being lower than two years, suggesting very rapid changes and extensive turnover of local climates in the PAs (Brito-Morales et al., 2018; Heikkinen et al., 2020; Nila & Hossain, 2019). PAs are expected to play a key role as stepping-stones for species dispersal to new regions (Gillingham & Thomas, 2023; Thomas & Gillingham, 2015), but species residing in small PAs within human-dominated flatlands may increasingly lose suitable conditions and face difficulties in shifting across space in pace with climate change (Hannah et al., 2014; Kharouba & Kerr, 2010; Parks et al., 2023).

### 5.3 Implications for developing a climate-wise Trans-European Nature Network

Our results suggest that climate change-based risks have largely been ignored in the designation of European PAs, potentially undermining their future conservation value. Simply designating 30% of Europe as “protected” does not guarantee that the biodiversity within these areas will persist in the long term, particularly if those areas are vulnerable to the adverse effects of climate change. The results of our and similar studies can be used to identify areas that are subject to more intensive climate change or a change via multiple dimensions of climate disruption. We identified such risks especially in the Boreal, Continental, and Steppic bioregions. In these areas a laissez-faire approach will likely not suffice, and specific interventions such as habitat restoration, creation of microhabitat, and translocations, including assisted migration (Hällfors et al., 2016) along velocity gradients, may be needed to maintain biodiversity.

High climate change pressures revealed by our study support the recommendations for integrating climate-wise considerations into nature conservation and management planning (Brito-Morales et al., 2018; Nadeau et al., 2015). This is of particular importance when planning the PA expansion in Europe, as part of a Trans-European Nature Network (Jung et al., 2024; Lai et al., 2022; Rannow & Förster, 2014). Many earlier recommendations remain valid also in climate-wise conservation planning, including: (i) determining key areas where species and habitats are most exposed to *vs* better buffered against climate change, (ii) targeting new conservation actions to increase the topographical heterogeneity of PAs and thereby supporting longer climate residence times and extending species holdouts, (iii) increasing landscape permeability and ecological connectivity in areas experiencing high climate velocities to enhance species movements, (iv) tailoring management and restoration in PAs to improve the resilience of species populations to climate change impacts, and (v) establishing new PAs in areas where climate exposure risk are low and the potential for climate refugia are high (Hannah et al., 2014; Mawdsley, 2011; Nadeau & Fuller, 2015; Thomas & Gillingham, 2015).

Priority setting for new conservation areas need to be carefully considered and account for several factors, e.g., species adaptive capacity, ecological importance, habitat fragmentation, habitat restoration, and connectivity potential. That said, other conditions being equal, it may be more strategic to establish new PAs in velocity coldspots as these can provide holdouts and climate refugia for species. Our results draw attention to the high velocity areas in the lowlands of Europe, highlighting the importance of increasing the permeability of the matrix areas and connectivity between existing PAs therein to allow species range shifting. However, areas with the biggest magnitude of climatic change diverge from those with the highest velocity. In these areas, in addition to enhancing connectivity, adaptative management of species populations and their habitats, combined with reducing pressures from other stressors should be prioritized. Ultimately, species conservation translocations may be considered as part of climate-wise conservation planning (Mawdsley, 2011; Thomas & Gillingham, 2015).

The failure to recognize and address the long-term impacts of climate change on the effectiveness of protected areas could lead to conservation efforts that are insufficient. A climate-proof Trans-European Nature Network should opt for spatial planning that simultaneously prioritizes climatically resilient areas, areas representing a variety of biomes and latitudes, and corridors that provide connectivity between these areas, to allow species the best available opportunities to thrive and move along with a changing climate.

## 6. Acknowledgements

This research was funded from the project NaturaConnect, which receives funding under the European Union’s Horizon Europe research and innovation programme under grant agreement number 101060429. Views and opinions expressed are however those of the authors only and do not necessarily reflect those of the European Union or the European Research Executive Agency. Neither the European Union nor the granting authority can be held responsible for them. MHH acknowledges funding by the Research Council of Finland (grant 360742), and MHH and RKH from the Finnish Strategic Research Council through the JUST TRANSITION programme for the project MUST (grant 358367).

## 7. Data availability

The data will be made available at https://zenodo.org/records/14146118

