## Supplementary Material for "The accelerating exposure of European protected areas to climate change"

### Appendix S1

### Climatic exposure for species within Natura2000 sites

We calculated the level of climate exposure for the species that occur in the Natura 2000 network using four climate metrics: local velocity, distance velocity, magnitude of change and residence time. In our analysis the aim was to improve the understanding of potential biodiversity implications of climate exposure affecting the Natura 2000 network. We used the Natura 2000 data provided by the European Environmental Agency (EEA,2023) to determine the association between the species and sites.

There are 2330 species included in the Natura 2000 list of species. Using this set of species as a starting point, we applied the Threat classification scheme of the IUCN Red List (IUCN 2023) to select species classified as threatened by climate change. This filtering provided us with a set of 1011 focal IUCN species that were then entered in the next step of our focal species selection process.

The 1011 IUCN species data were next cross-checked with the list of European species located inside Natura 2000 sites to create a new list containing only species present in Europe that have already been identified as vulnerable to climate change. We accounted for potential misspellings or changes of scientific names using the Global Name Verifier (Global Names Verifier, 2023), a software that allows checking a list of species against the main databases of biodiversity, assigning a score to each matched name. Fuzzy matching, a feature that accounts for potential misspellings of names, was employed in this step. We acknowledge that fuzzy matching is a function which needs to be used with some caution. Indeed, some studies report (Patterson et al., 2016) that the method can produce errors due to confusing similar species’ names. Here, we accounted for this potential error by limiting the number of characters changed to 1 and manually checking for species with more than 1 character changed.

### Defining variables used:

All four climate exposure metrics calculated for the two key variables: “annual mean temperature” and ”total annual precipitation”. When these two variables were used in combination, we refer to this summarised variable as “climate”.

### **Table S1** Metrics and their combination calculated for the whole Europe.

| **Metrics** | **ESMs** | **SSP-RCP Scenarios** | **Ensemble** |
| --- | --- | --- | --- |
| Local velocity | GFDL | SSP 1-2.6 | ✓ |
|  | MPI | SSP 3-7.0 |  |
|  | UKESM | SSP 5-8.5 |  |
| Distance velocity | GFDL | SSP 1-2.6 | ✓ |
|  | MPI | SSP 3-7.0 |  |
|  | UKESM | SSP 5-8.5 |  |
| Magnitude | GFDL | SSP 1-2.6 | ✓ |
|  | MPI | SSP 3-7.0 |  |
|  | UKESM | SSP 5-8.5 |  |
| Residence time | Calculated only on the ensemble of the local velocity, for PAs | SSP 1-2.6 | X |
|  |  | SSP 3-7.0 |  |
|  |  | SSP 5-8.5 |  |

### **Table S2** Variables used for propensity score matching, with their description, unit of measure and relative source.

| **Variable** | **Description** | **Unit** | **Description** | **Source** |
| --- | --- | --- | --- | --- |
| bio1 | mean annual air temperature | °C | mean annual daily mean air temperatures averaged over 1 year | Chelsa V 2.1,(Karger et al., 2017, 2021) |
| bio2 | mean diurnal air temperature range | °C | mean diurnal range of temperatures averaged over 1 year | Chelsa V 2.1,(Karger et al., 2017, 2021) |
| bio3 | isothermality | °C | ratio of diurnal variation to annual variation in temperatures | Chelsa V 2.1,(Karger et al., 2017, 2021) |
| bio4 | temperature seasonality | °C/100 | standard deviation of the monthly mean temperatures | Chelsa V 2.1,(Karger et al., 2017, 2021) |
| bio5 | mean daily maximum air temperature of the warmest month | °C | The highest temperature of any monthly daily mean maximum temperature | Chelsa V 2.1,(Karger et al., 2017, 2021) |
| bio6 | mean daily minimum air temperature of the coldest month | °C | The lowest temperature of any monthly daily mean maximum temperature | Chelsa V 2.1,(Karger et al., 2017, 2021) |
| bio7 | annual range of air temperature | °C | The difference between the Maximum Temperature of Warmest month and the Minimum Temperature of Coldest month | Chelsa V 2.1,(Karger et al., 2017, 2021) |
| bio8 | mean daily mean air temperatures of the wettest quarter | °C | The wettest quarter of the year is determined (to the nearest month) | Chelsa V 2.1,(Karger et al., 2017, 2021) |
| bio9 | mean daily mean air temperatures of the driest quarter | °C | The driest quarter of the year is determined (to the nearest month) | Chelsa V 2.1,(Karger et al., 2017, 2021) |
| bio10 | mean daily mean air temperatures of the warmest quarter | °C | The warmest quarter of the year is determined (to the nearest month) | Chelsa V 2.1,(Karger et al., 2017, 2021) |
| bio11 | mean daily mean air temperatures of the coldest quarter | °C | The coldest quarter of the year is determined (to the nearest month) | Chelsa V 2.1,(Karger et al., 2017, 2021) |
| bio12 | annual precipitation amount | kg m^-2^ | Accumulated precipitation amount over 1 year | Chelsa V 2.1,(Karger et al., 2017, 2021) |
| bio13 | precipitation amount of the wettest month | kg m^-2^ | The precipitation of the wettest month. | Chelsa V 2.1,(Karger et al., 2017, 2021) |
| bio14 | precipitation amount of the driest month | kg m^-2^ | The precipitation of the driest month. | Chelsa V 2.1,(Karger et al., 2017, 2021) |
| bio15 | precipitation seasonality | kg m^-2^ | The Coefficient of Variation is the standard deviation of the monthly precipitation estimates expressed as a percentage of the mean of those estimates (i.e. the annual mean) | Chelsa V 2.1,(Karger et al., 2017, 2021) |
| bio16 | mean monthly precipitation amount of the wettest quarter | kg m^-2^ | The wettest quarter of the year is determined (to the nearest month) | Chelsa V 2.1,(Karger et al., 2017, 2021) |
| bio17 | mean monthly precipitation amount of the driest quarter | kg m^-2^ | The driest quarter of the year is determined (to the nearest month) | Chelsa V 2.1,(Karger et al., 2017, 2021) |
| bio18 | mean monthly precipitation amount of the warmest quarter | kg m^-2^ | The warmest quarter of the year is determined (to the nearest month) | Chelsa V 2.1,(Karger et al., 2017, 2021) |
| bio19 | mean monthly precipitation amount of the coldest quarter | kg m^-2^ | The coldest quarter of the year is determined (to the nearest month) | Chelsa V 2.1,(Karger et al., 2017, 2021) |
| Landcover fraction | The classes used were:  bare and sparse vegetation, forest,seasonal inland water, permanent inland water, built-up, herbaceous vegetation, shrubland, cropland, moss and lichen, snow and ice. | % | Landcover fraction refers to the proportion of a specific land cover type (e.g., forest, urban, inland water) within a given area. It quantifies the relative contribution of different land cover categories to the total land area. | Copernicus (Buchhorn et al., 2020) |
| Human footprint |  | Composite index (1 to 50) | Human footprint measures the extent of human influence or impact on a area, considering factors such as infrastructure, population density, and land use | Venter et al., 2016 |
| Elevation |  | m | It refers to the height of a point on the Earth's surface above sea level. It is a crucial topographic characteristic influencing climate, ecosystems, and human activities. | Worldclim (Fick & Hijmans, 2017) |
| Slope |  | Degrees (°) | It represents the steepness or incline of the terrain at a specific location. | Terra: terrainr function (calculated from elevation data) (Hijmans, 2023) |
| Aspect |  | Degrees (°) | describes the compass direction that a slope faces. It is measured in degrees clockwise from north and provides information about the orientation of a land surface. | Terra: terrainr function (calculated from elevation data)(Hijmans, 2023) |
| Roughness |  |  | It refers to the irregularity or variability of the Earth's surface. It can be characterized by variations in elevation, land cover, or other surface features. | *Terra*: *terrainr* function (calculated from elevation data) (Hijmans, 2023) |

### Table S3 Median metrics values for every biogeographic region. Velocity are reported in km/y while magnitude is unitless.

| **Bioregion** | **Local SSP1-2.6** | **Local SSP3-7.0** | **Local SSP5-8.5** | **Distance SSP1-2.6** | **Distance SSP3-7.0** | **Distance SSP5-8.5** | **Magnitude SSP1-2.6** | **Magnitude SSP3-7.0** | **Magnitude SSP5-8.5** |
| --- | --- | --- | --- | --- | --- | --- | --- | --- | --- |
| Alpine | 0.087 | 0.204 | 0.322 | 0.069 | 0.154 | 0.209 | 4.91 | 6.06 | 7.39 |
| Arctic | 0.178 | 0.457 | 0.570 | 0.067 | 0.272 | 0.264 | 4.82 | 5.89 | 6.93 |
| Atlantic | 0.521 | 0.738 | 1.173 | 0.313 | 1.190 | 1.515 | 3.96 | 4.80 | 6.30 |
| Black Sea | 0.912 | 1.369 | 2.101 | 0.440 | 0.826 | 1.096 | 3.84 | 4.96 | 6.03 |
| Boreal | 0.915 | 1.769 | 3.055 | 1.617 | 2.268 | 2.039 | 4.13 | 5.13 | 6.58 |
| Continental | 0.855 | 1.450 | 2.284 | 0.622 | 0.964 | 1.200 | 3.97 | 4.84 | 6.44 |
| Macaronesian | 0.234 | 0.217 | 0.342 | 0.039 | 0.066 | 0.078 | 2.74 | 3.68 | 4.40 |
| Mediterranean | 0.163 | 0.414 | 0.656 | 0.105 | 0.197 | 0.232 | 4.79 | 6.36 | 7.19 |
| Pannonian | 2.160 | 4.850 | 7.161 | 0.536 | 0.869 | 1.047 | 3.53 | 4.57 | 5.84 |
| Steppic | 1.103 | 2.361 | 3.518 | 1.531 | 2.754 | 2.840 | 3.07 | 3.80 | 5.12 |

### **Table S4** C-statistic before and after matching for every biomes.

| **Biomes** | **C stat pre matching** | **C stat post matching** |
| --- | --- | --- |
| Alpine | 0.745 | 0.516 |
| Arctic | 0.745 | 0.509 |
| Atlantic | 0.745 | 0.508 |
| Boreal | 0.745 | 0.635 |
| Continental | 0.745 | 0.500 |
| Mediterranean | 0.745 | 0.518 |
| Pannonian | 0.745 | 0.595 |
| Steppic | 0.745 | 0.517 |
| BlackSea | 0.745 | 0.524 |
| Macaronesia | 0.745 | 0.532 |

### Table S5 Wilcoxon's test results after propensity score matching reaggregated on the European level. Mean and median values for inside PAs (1) and outside control areas (0) for Annual Mean Temperature. Total Annual Precipitation and Climate. Velocity values are reported in km/y. Significance codes: P<0.001: ***, P<0.01: **, P<0.05: *.

| **Protection** | **Metric** | **Scenario** | **Variable** | **Metric value** | **W/P.value** | **Value** | **Significance code** |
| --- | --- | --- | --- | --- | --- | --- | --- |
| 0 | local | hist | Climate | 1.498 | W | 7.724E+11 |  |
| 1 | local | hist | Climate | 1.756 | P | 0 | *** |
| 0 | local | SSP1-2.6 | Climate | 0.976 | W | 7.678E+11 |  |
| 1 | local | SSP1-2.6 | Climate | 1.184 | P | 0 | *** |
| 0 | local | SSP3-7.0 | Climate | 1.787 | W | 7.754E+11 |  |
| 1 | local | SSP3-7.0 | Climate | 2.124 | P | 2.55E-247 | *** |
| 0 | local | SSP5-8.5 | Climate | 2.771 | W | 7.804E+11 |  |
| 1 | local | SSP5-8.5 | Climate | 3.324 | P | 7.23E-138 | *** |
| 0 | local | hist | Total Annual Precipitation | 0.448 | W | 7.674E+11 |  |
| 1 | local | hist | Total Annual Precipitation | 0.542 | P | 0 | *** |
| 0 | local | SSP1-2.6 | Total Annual Precipitation | 0.207 | W | 7.281E+11 |  |
| 1 | local | SSP1-2.6 | Total Annual Precipitation | 0.275 | P | 0 | *** |
| 0 | local | SSP3-7.0 | Total Annual Precipitation | -0.172 | W | 7.582E+11 |  |
| 1 | local | SSP3-7.0 | Total Annual Precipitation | -0.141 | P | 0 | *** |
| 0 | local | SSP5-8.5 | Total Annual Precipitation | -0.290 | W | 8.039E+11 |  |
| 1 | local | SSP5-8.5 | Total Annual Precipitation | -0.318 | P | 6.941E-55 | *** |
| 0 | local | hist | Annual Mean Temperature | 0.935 | W | 7.767E+11 |  |
| 1 | local | hist | Annual Mean Temperature | 1.106 | P | 1.58E-217 | *** |
| 0 | local | SSP1-2.6 | Annual Mean Temperature | 0.629 | W | 7.969E+11 |  |
| 1 | local | SSP1-2.6 | Annual Mean Temperature | 0.784 | P | 0.0005691 | *** |
| 0 | local | SSP3-7.0 | Annual Mean Temperature | 1.500 | W | 7.788E+11 |  |
| 1 | local | SSP3-7.0 | Annual Mean Temperature | 1.846 | P | 9.75E-170 | *** |
| 0 | local | SSP5-8.5 | Annual Mean Temperature | 2.363 | W | 7.781E+11 |  |
| 1 | local | SSP5-8.5 | Annual Mean Temperature | 2.897 | P | 4.58E-184 | *** |
| 0 | distance | SSP1-2.6 | Climate | 0.156 | W | 5.56E+11 |  |
| 1 | distance | SSP1-2.6 | Climate | 0.175 | P | 2.64E-07 | *** |
| 0 | distance | SSP3-7.0 | Climate | 0.366 | W | 5.43E+11 |  |
| 1 | distance | SSP3-7.0 | Climate | 0.455 | P | 1.13E-114 | *** |
| 0 | distance | SSP5-8.5 | Climate | 0.462 | W | 5.33E+11 |  |
| 1 | distance | SSP5-8.5 | Climate | 0.610 | P | 0.00E+00 | *** |
| 0 | magnitude | SSP1-2.6 | Climate | 4.427 | W | 8.28E+11 |  |
| 1 | magnitude | SSP1-2.6 | Climate | 4.360 | P | 0 | *** |
| 0 | magnitude | SSP3-7.0 | Climate | 5.565 | W | 8.354E+11 |  |
| 1 | magnitude | SSP3-7.0 | Climate | 5.466 | P | 0 | *** |
| 0 | magnitude | SSP5-8.5 | Climate | 6.903 | W | 8.506E+11 |  |
| 1 | magnitude | SSP5-8.5 | Climate | 6.774 | P | 0 | *** |
| 0 | magnitude | SSP1-2.6 | Total Annual Precipitation | 2.188 | W | 7.999E+11 |  |
| 1 | magnitude | SSP1-2.6 | Total Annual Precipitation | 2.177 | P | 1.325E-18 | *** |
| 0 | magnitude | SSP3-7.0 | Total Annual Precipitation | 2.253 | W | 7.999E+11 |  |
| 1 | magnitude | SSP3-7.0 | Total Annual Precipitation | 2.245 | P | 3.536E-18 | *** |
| 0 | magnitude | SSP5-8.5 | Total Annual Precipitation | 3.954 | W | 8.11E+11 |  |
| 1 | magnitude | SSP5-8.5 | Total Annual Precipitation | 3.913 | P | 7.5E-171 | *** |
| 0 | magnitude | SSP1-2.6 | Annual Mean Temperature | 3.440 | W | 8.217E+11 |  |
| 1 | magnitude | SSP1-2.6 | Annual Mean Temperature | 3.377 | P | 0 | *** |
| 0 | magnitude | SSP3-7.0 | Annual Mean Temperature | 4.749 | W | 8.295E+11 |  |
| 1 | magnitude | SSP3-7.0 | Annual Mean Temperature | 4.652 | P | 0 | *** |
| 0 | magnitude | SSP5-8.5 | Annual Mean Temperature | 5.178 | W | 8.35E+11 |  |
| 1 | magnitude | SSP5-8.5 | Annual Mean Temperature | 5.062 | P | 0 | *** |

### Table S6 Wilcoxon's test results after propensity score matching on the biomes level. Mean and median values of local velocity and magnitude for inside PAs (1) and outside control areas (0) for mean Annual Mean Temperature, Total Annual Precipitation and Climate. Velocity values are reported in km/y, magnitude is adimensional. Significance codes: P<0.001: ***, P<0.01: **, P<0.05: *.

**See the excel table S6**

### Table S7 Projected mean and median climate residence time (years) under different SSPs, for each biome in Europe.

| **Biome** | **Mean resT SSP1-2.6sum** | **Mean resT SSP3-7.0sum** | **Mean resT SSP5-8.5sum** | **Median resT SSP1-2.6sum** | **Median resT SSP3-7.0sum** | **Median resT SSP5-8.5sum** |
| --- | --- | --- | --- | --- | --- | --- |
| Alpine | 84.6 | 45.0 | 30.9 | 41.0 | 24.2 | 16.8 |
| Arctic | 90.1 | 32.6 | 25.1 | 67.6 | 14.4 | 13.5 |
| Atlantic | 24.2 | 10.6 | 6.9 | 3.1 | 2.3 | 1.4 |
| BlackSea | 8.3 | 5.3 | 3.4 | 3.6 | 2.5 | 1.5 |
| Boreal | 5.1 | 2.2 | 1.4 | 1.9 | 1.1 | 0.6 |
| Continental | 9.8 | 6.4 | 4.0 | 4.0 | 2.6 | 1.7 |
| Macaronesia | 29.2 | 18.0 | 17.3 | 14.2 | 12.5 | 10.3 |
| Mediterranean | 57.6 | 21.0 | 13.0 | 26.7 | 11.8 | 7.1 |
| Pannonian | 6.8 | 2.9 | 2.0 | 2.2 | 1.0 | 0.7 |
| Steppic | 6.8 | 4.4 | 2.5 | 3.2 | 1.6 | 1.1 |

### Table S8 Median climate metrics values for all species threatened by climate change occurring within Natura 2000 (1011 species), separated by taxonomic group, under scenarios SSP1-2.6, SSP3-7.0, SSP5-8.5 . Velocity values are reported in km/y.

|  | **Local sum hist median** | **Local sum ssp1-2.6 median** | **Local sum ssp3-7.0 median** | **Local sum ssp5-8.5 median** | **Magnitude sum ssp1-2.6 median** | **Magnitude sum ssp3-7.0 median** | **Magnitude sum ssp5-8.5 median** |
| --- | --- | --- | --- | --- | --- | --- | --- |
| All species | 1.701 | 1.178 | 2.094 | 3.451 | 4.235 | 5.304 | 6.659 |
| Amphibians | 1.691 | 1.141 | 2.042 | 3.274 | 4.341 | 5.340 | 6.775 |
| Birds | 2.099 | 1.433 | 2.463 | 3.978 | 4.221 | 5.271 | 6.645 |
| Invertebrates | 0.961 | 0.557 | 1.401 | 2.135 | 4.278 | 5.439 | 6.688 |
| Mammals | 2.237 | 1.266 | 2.116 | 3.573 | 4.245 | 5.329 | 6.659 |
| Plants | 1.085 | 0.684 | 1.450 | 2.438 | 4.177 | 5.254 | 6.628 |
| Reptiles | 1.468 | 0.959 | 1.432 | 2.511 | 4.351 | 5.442 | 6.776 |

### Table S9 Median climate metrics values for the 514 species with spatial distribution information available from the IUCN Red List divided by main taxonomic group, according to their protected range under scenarios SSP1-2.6, SSP3-7.0, SSP5-8.5 . Velocity values are reported in km/y.

|  | **Cover fraction** | **Local sum hist** | **Local sum SSP1-2.6** | **Local sum SSP3-7.0** | **Local sum SSP5-8.5** | **Magnitude sum SSP1-2.6** | **Magnitude sum SSP3-7.0** | **Magnitude sum SSP5-8.5** |
| --- | --- | --- | --- | --- | --- | --- | --- | --- |
| All species | 0-25 | 1.973 | 1.311 | 2.292 | 3.768 | 4.228 | 5.299 | 6.656 |
|  | 25-50 | 1.373 | 0.847 | 1.583 | 2.687 | 4.242 | 5.331 | 6.659 |
|  | 50-75 | 0.848 | 0.393 | 0.638 | 1.294 | 4.153 | 5.26 | 6.61 |
|  | 75-100 | 0.728 | 0.373 | 0.597 | 1.169 | 4.339 | 5.457 | 6.806 |
| Amphibians | 0-25 | 1.686 | 1.138 | 1.939 | 3.237 | 4.354 | 5.508 | 6.826 |
|  | 25-50 | 1.696 | 1.222 | 2.145 | 3.593 | 4.18 | 5.255 | 6.652 |
|  | 50-75 | 3.325 | 1.992 | 4.759 | 6.68 | 4.341 | 5.394 | 6.715 |
|  | 75-100 | 0.132 | 0.357 | 0.46 | 0.627 | 4.772 | 6.524 | 7.21 |
| Birds | 0-25 | 2.112 | 1.453 | 2.472 | 4.009 | 4.215 | 5.263 | 6.642 |
|  | 25-50 | 1.748 | 1.269 | 2.385 | 3.728 | 4.247 | 5.358 | 6.805 |
|  | 75-100 | 0.776 | 0.423 | 0.94 | 1.392 | 4.29 | 5.269 | 6.807 |
| Invertebrates | 0-25 | 1.455 | 0.913 | 1.791 | 3.249 | 4.231 | 5.342 | 6.667 |
|  | 25-50 | 0.539 | 0.418 | 0.835 | 1.531 | 4.433 | 5.597 | 6.832 |
|  | 50-75 | 0.522 | 0.26 | 0.561 | 0.743 | 4.403 | 5.767 | 6.739 |
|  | 75-100 | 0.961 | 0.393 | 1.007 | 1.511 | 4.467 | 5.713 | 7.008 |
| Mammals | 0-25 | 2.471 | 1.359 | 2.266 | 3.783 | 4.25 | 5.31 | 6.67 |
|  | 25-50 | 1.576 | 1.099 | 1.882 | 3.335 | 4.245 | 5.331 | 6.612 |
|  | 75-100 | 1.44 | 0.932 | 1.438 | 2.446 | 4.235 | 5.423 | 6.756 |
| Plants | 0-25 | 1.467 | 1.002 | 1.723 | 3.014 | 4.204 | 5.266 | 6.643 |
|  | 25-50 | 1.073 | 0.667 | 1.412 | 2.267 | 4.17 | 5.253 | 6.634 |
|  | 50-75 | 0.95 | 0.408 | 0.912 | 1.599 | 4.057 | 5.254 | 6.557 |
|  | 75-100 | 0.326 | 0.224 | 0.588 | 0.783 | 4.396 | 5.605 | 6.777 |
| Reptiles | 0-25 | 1.633 | 1.002 | 2.22 | 3.204 | 4.766 | 5.84 | 7.376 |
|  | 25-50 | 1.353 | 0.842 | 1.09 | 1.789 | 4.241 | 5.388 | 6.717 |
|  | 50-75 | 1.58 | 1.293 | 2.831 | 3.851 | 4.858 | 6.19 | 7.4 |
|  | 75-100 | 2.544 | 1.782 | 2.557 | 3.893 | 4.235 | 5.357 | 6.704 |

### Table S10 Weighted mean climate metrics for species with more than 50% of their European terrestrial range located inside the Natura 2000 network. Velocity values are reported in km/y.

| **Species name** | **Local sum hist** | **Local sum SSP1-2.6** | **Local sum SSP3-7.0** | **Local sum SSP5-8.5** | **Magnitude sum SSP1-2.6** | **Magnitude sum SSP3-7.0** | **Magnitude sum SSP5-8.5** | **Red list category** | **Area km** | **Area km**  **Overlaps** | **Cover fraction** | **Group** |
| --- | --- | --- | --- | --- | --- | --- | --- | --- | --- | --- | --- | --- |
| *Calotriton arnoldi* | 0.132 | 0.357 | 0.460 | 0.627 | 4.772 | 6.524 | 7.210 | Critically Endangered | 289.5 | 289.5 | 100 | Amphibians |
| *Chalcides simonyi* | 3.030 | 2.275 | 4.850 | 6.768 | 4.495 | 5.628 | 7.050 | Endangered | 530.6 | 288.8 | 54.44 | Reptiles |
| *Crocidura zimmermanni* | 1.382 | 0.646 | 0.860 | 1.718 | 4.274 | 5.513 | 6.872 | Endangered | 900.2 | 851.5 | 94.59 | Mammals |
| *Fringilla teydea* | 0.481 | 0.439 | 0.940 | 1.392 | 4.290 | 5.269 | 6.807 | Near Threatened | 349.9 | 317.2 | 90.66 | Birds |
| *Iberolacerta aurelioi* | 0.131 | 0.311 | 0.811 | 0.933 | 5.221 | 6.751 | 7.750 | Endangered | 122.4 | 81.6 | 66.65 | Reptiles |
| *Iberolacerta bonnali* | 2.544 | 1.782 | 2.557 | 3.893 | 4.235 | 5.357 | 6.704 | Near Threatened | 1583.1 | 1227.2 | 77.52 | Reptiles |
| *Lyciasalamandra helverseni* | 0.375 | 0.169 | 0.206 | 0.308 | 5.539 | 6.841 | 8.150 | Vulnerable | 367.1 | 199.6 | 54.36 | Amphibians |
| *Lynx pardinus* | 1.498 | 1.218 | 2.016 | 3.175 | 4.197 | 5.332 | 6.640 | Endangered | 1191.7 | 909.6 | 76.33 | Mammals |
| *Pterodroma madeira* | 0.848 | 0.423 | 2.029 | 2.958 | 3.826 | 5.252 | 6.048 | Endangered | 363.7 | 346.1 | 95.18 | Birds |
| *Rana pyrenaica* | 3.325 | 2.394 | 4.935 | 7.321 | 4.341 | 5.394 | 6.715 | Endangered | 2237 | 1315.1 | 58.79 | Amphibians |
| *Speleomantes supramontis* | 4.249 | 1.992 | 4.759 | 6.680 | 3.820 | 5.096 | 6.045 | Endangered | 573.7 | 401.7 | 70.02 | Amphibians |

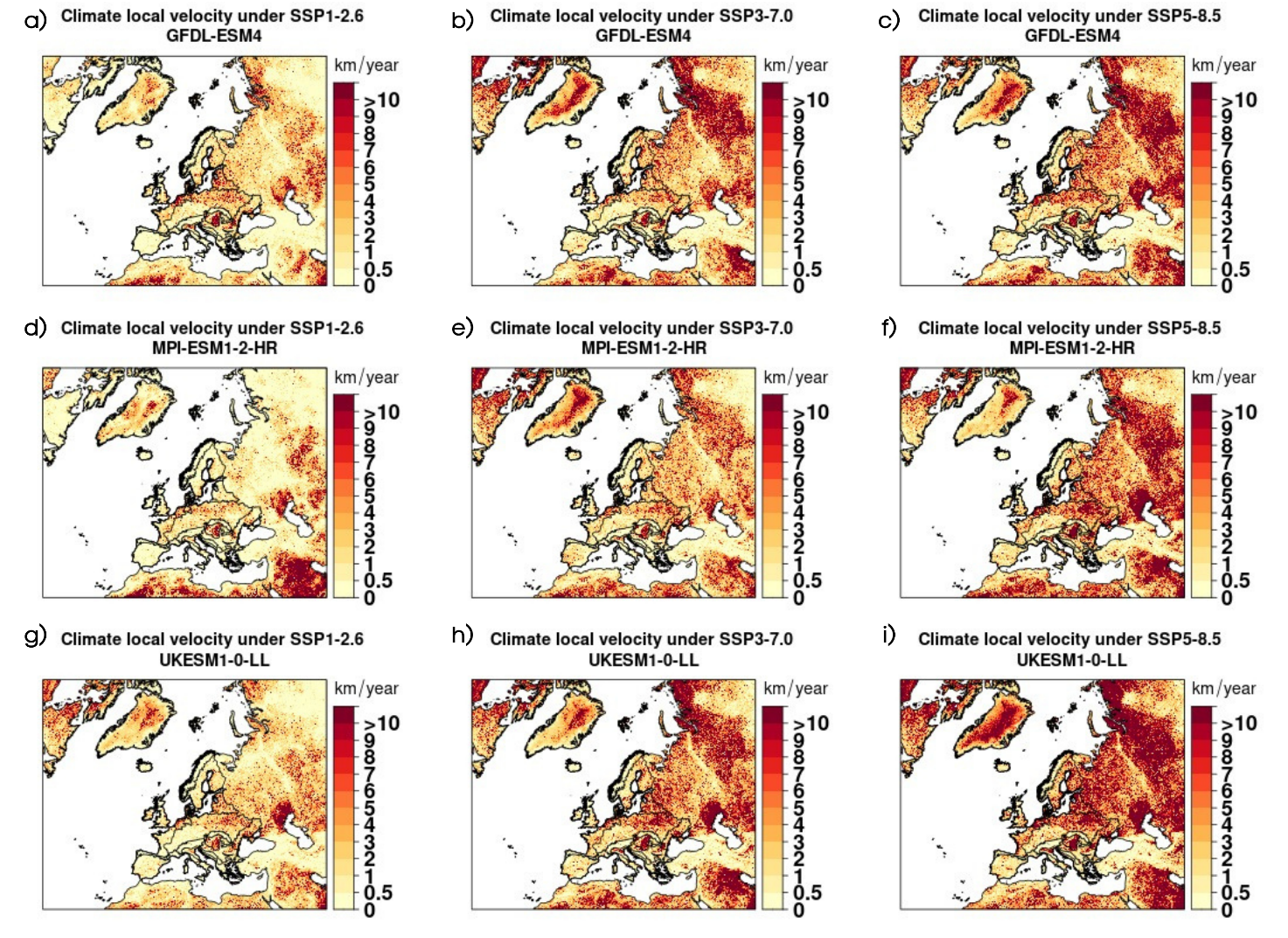

### Fig S1) Projections of change in local velocity under different ESMs and SSP-RCP scenariosfor local velocity of climate.

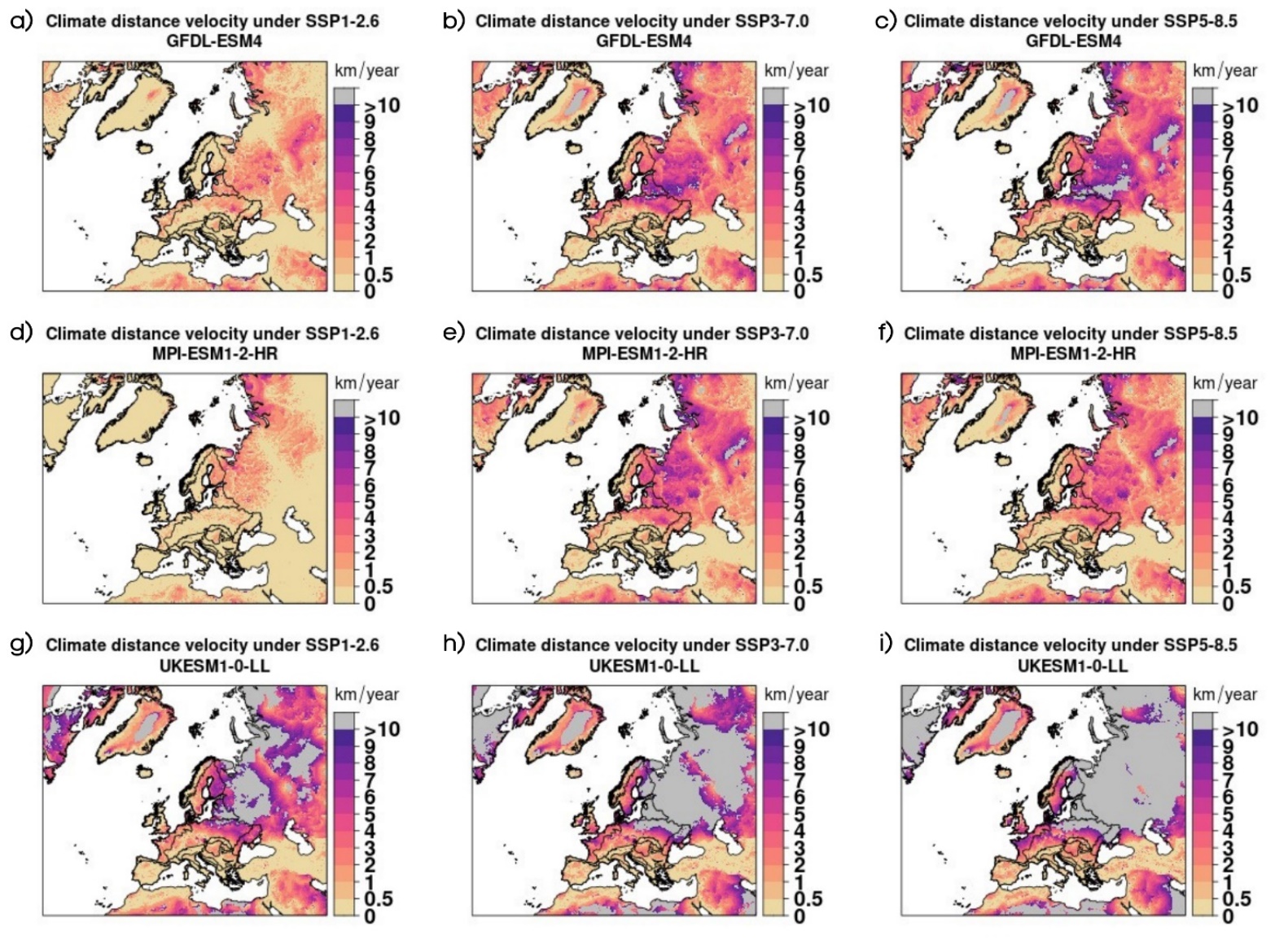

Fig S2) Projections of change in distance velocity under different ESMs and SSP-RCP scenarios. Area in grey are visible in multiple panels (particularly evident in c), g), h), and i)), indicating cells for which we found no climate analogue within 500 km.

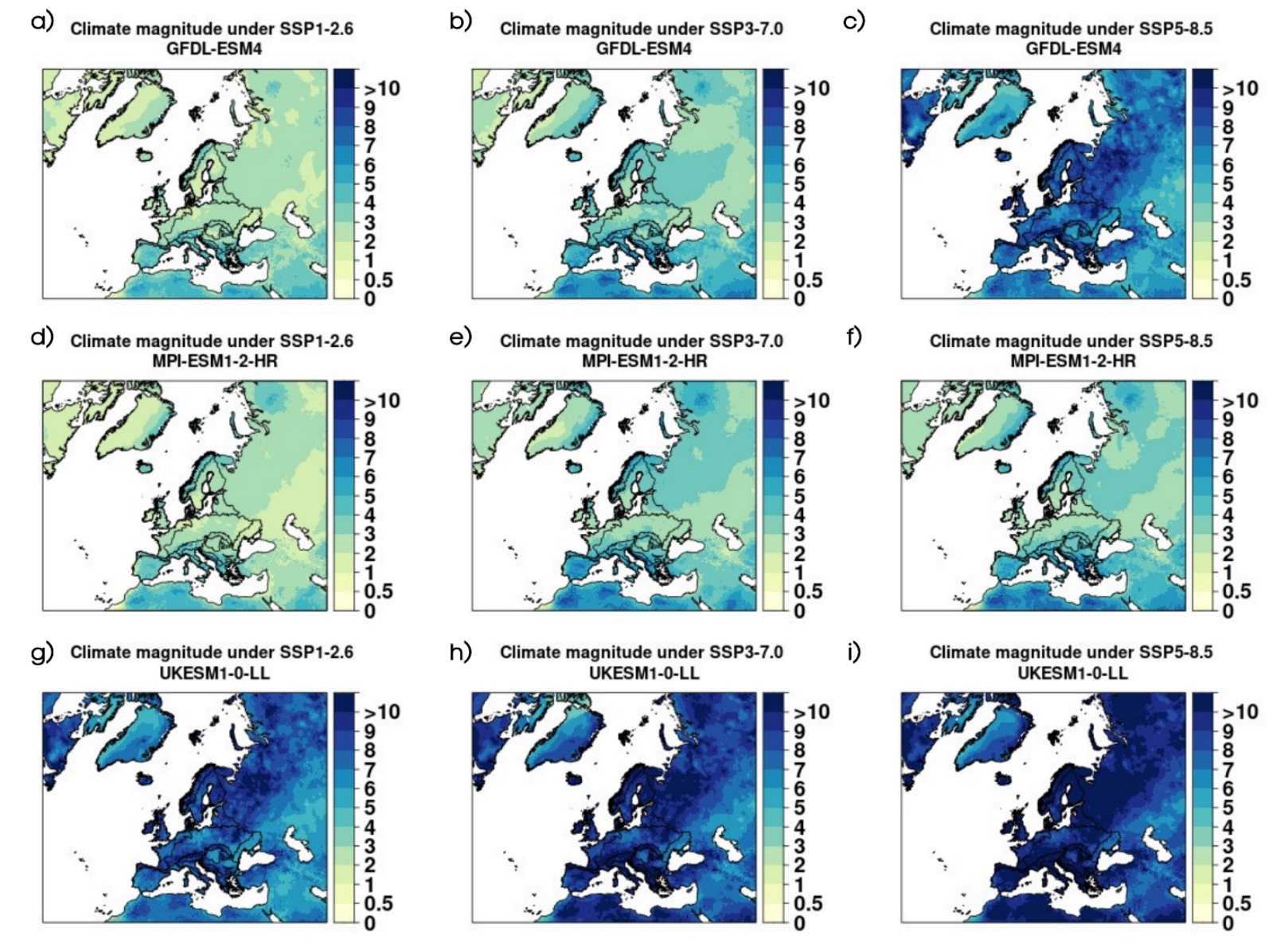

### Fig S**3**) Projections of change in magnitude under different ESMs and SSP-RCP scenarios for climate magnitude.

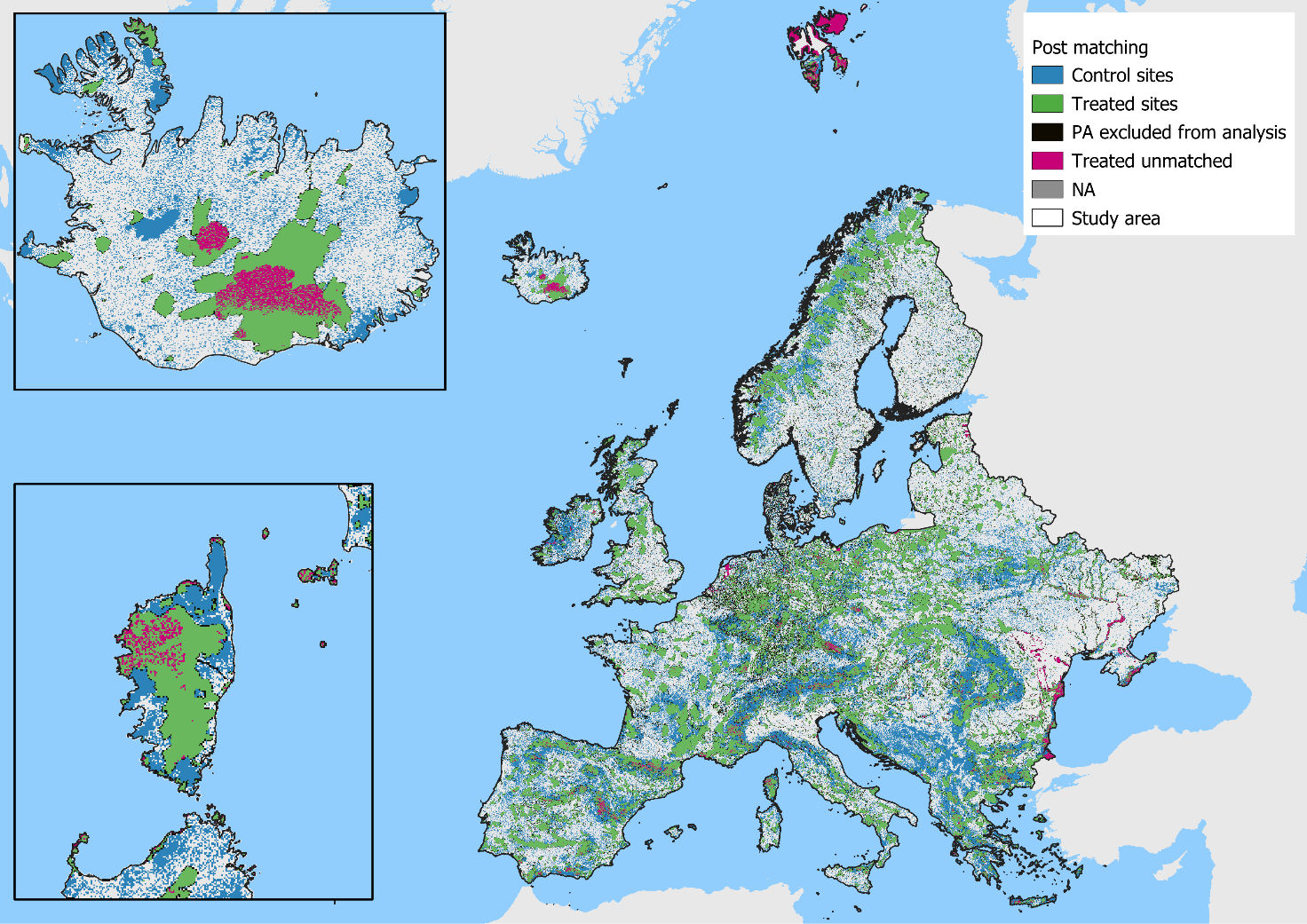

### Fig S4) Matching results: control sites which received a match are shown in green, unprotected matched cell (control) are shown in blue. Cells with PA coverage between 5 and 50% (excluded from the analysis) are shown in black. PAs for which we did not find a match ar shown in magenta. Unprotected areas nt selcted as control sites are shown in gray.

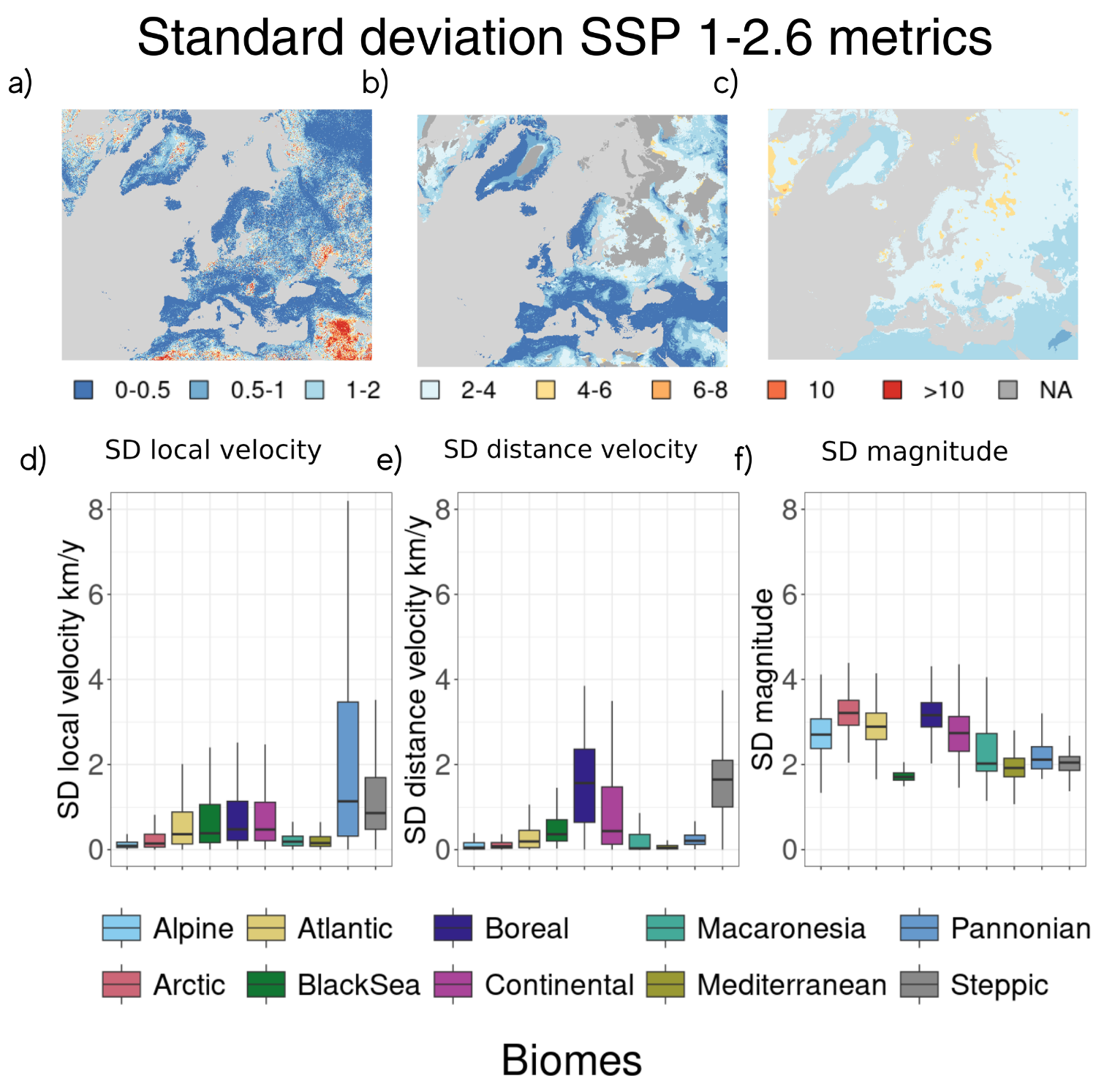

### Fig. S5) Projections of standard deviation for climate metrics, a) local velocity, b) distance velocity, c) magnitude under scenario SSP1-2.6 for the whole study area and their respective boxplots (d) ,e) and f)) reporting metrics value for each biogeographical region. For b) NA area represents cells for which one or more ESM found no analog climate within 500 km. In e) results are shown excluding NA cells.

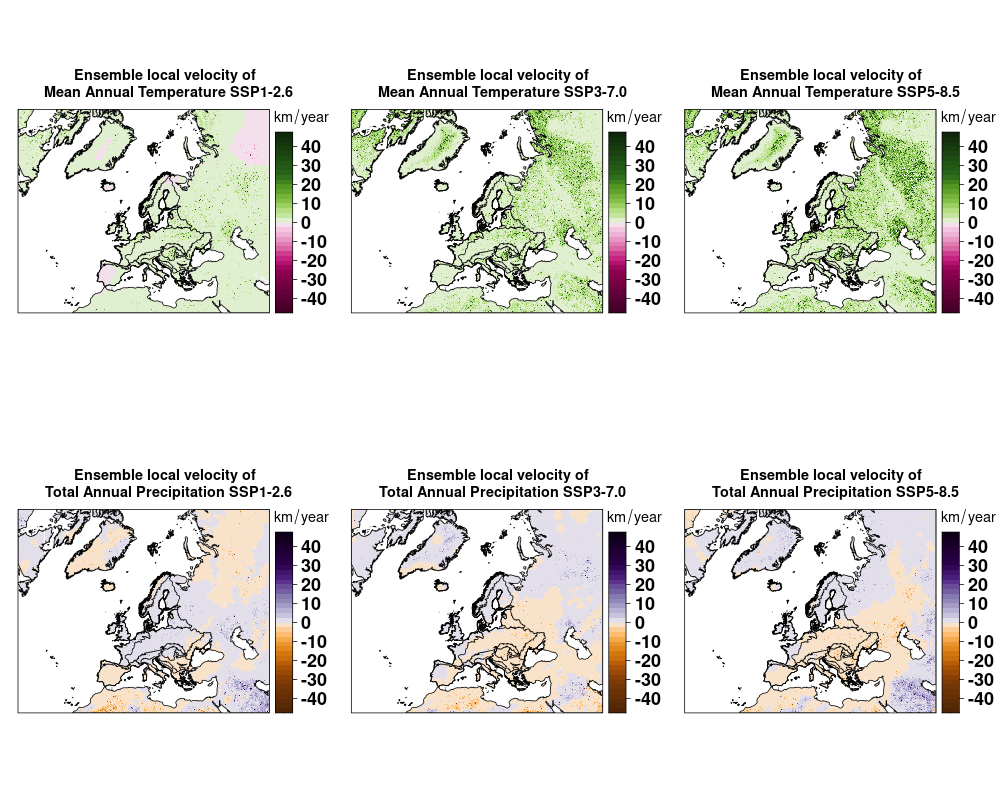

### Fig. S6) Projections of change in local velocity of temperature and precipitation under scenarios SSP1-2.6, SSP3-7.0, and SSP5-8.5..

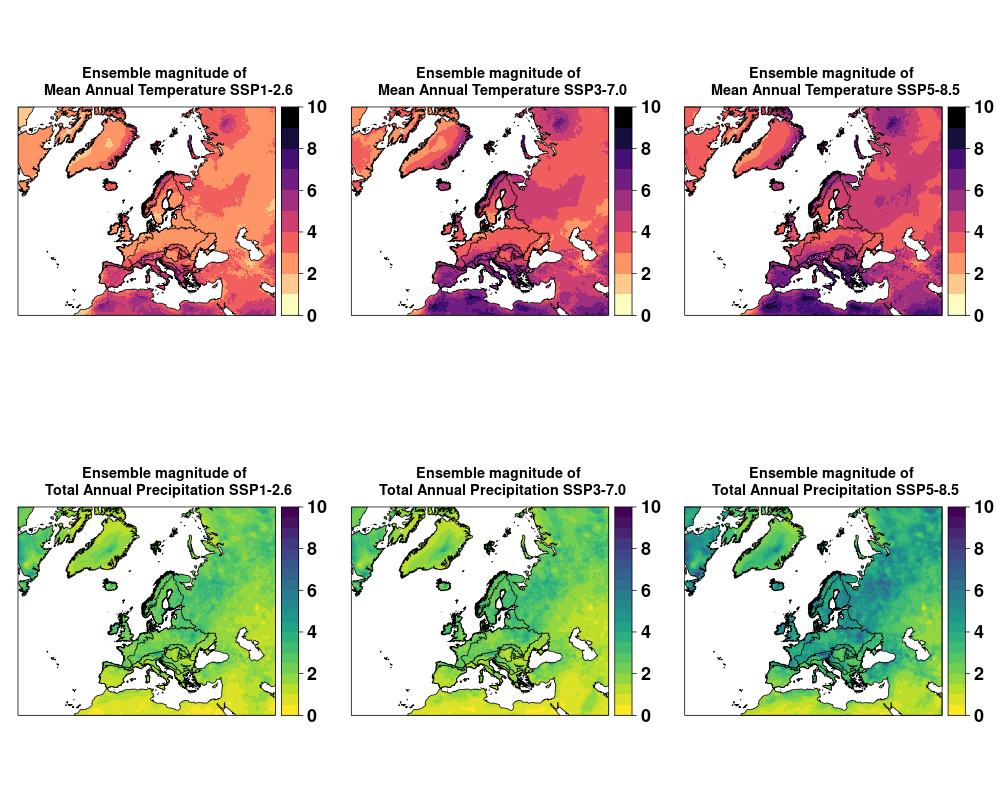

### Fig. S7) Projections of change magnitude of temperature and precipitation under scenarios SSP1-2.6, SSP3-7.0, and SSP5-8.5.

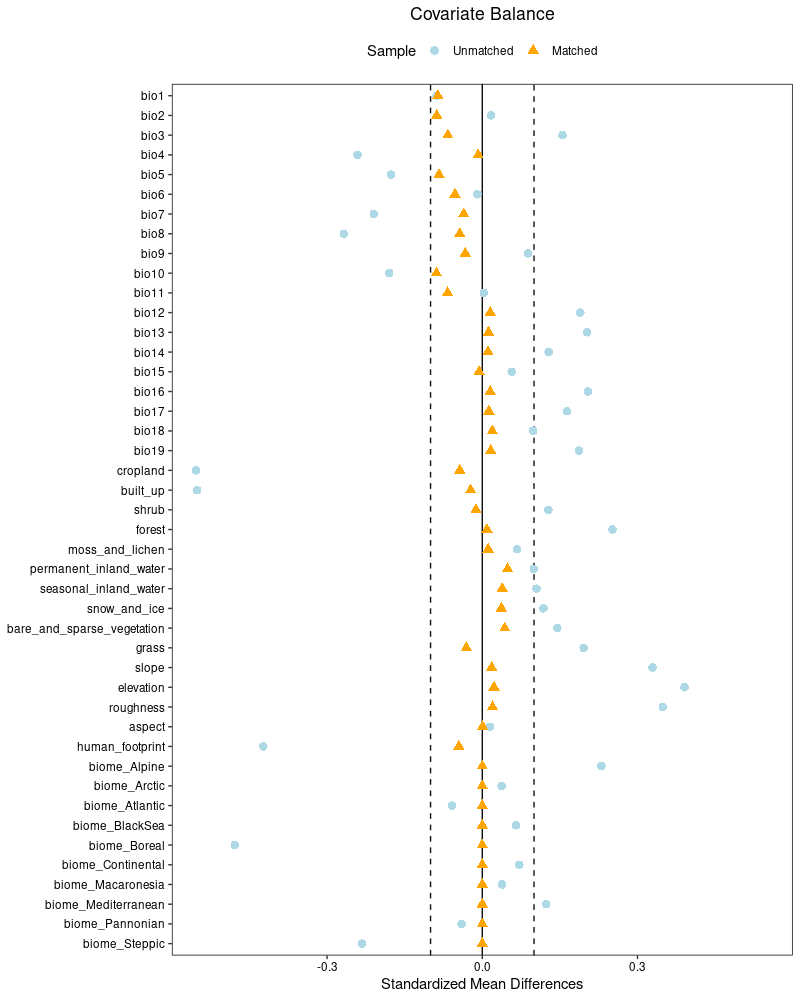

### Fig. S8) Love plot of the SMD test for the matching analysis of PAs with control sites.

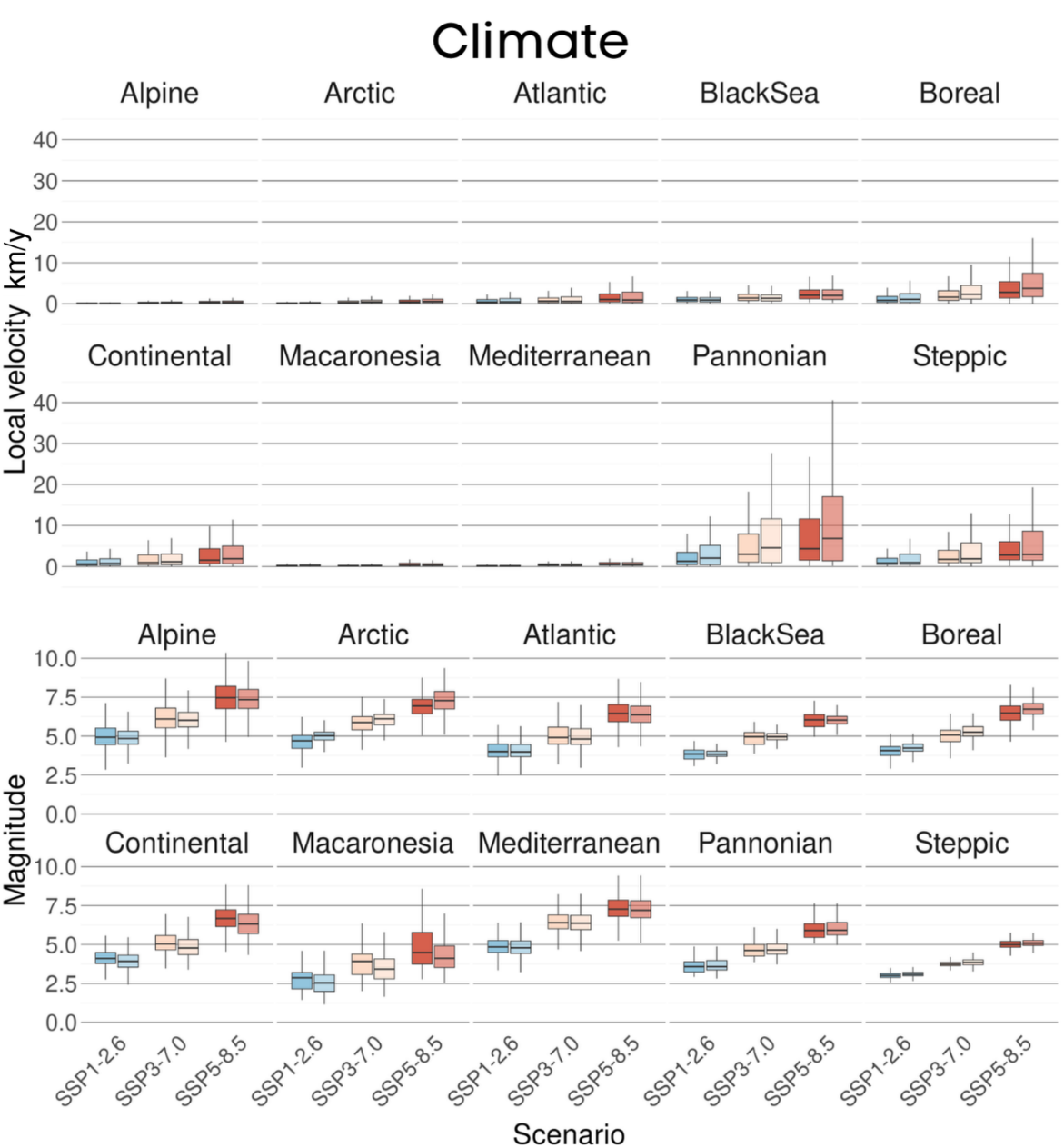

### Fig. S9) Boxplot of climate for local velocity, distance velocity, and magnitude in unprotected areas (darker colour) and PAs (lighter colour). Separate plots are reported for the 10 biomes covering the whole study area.

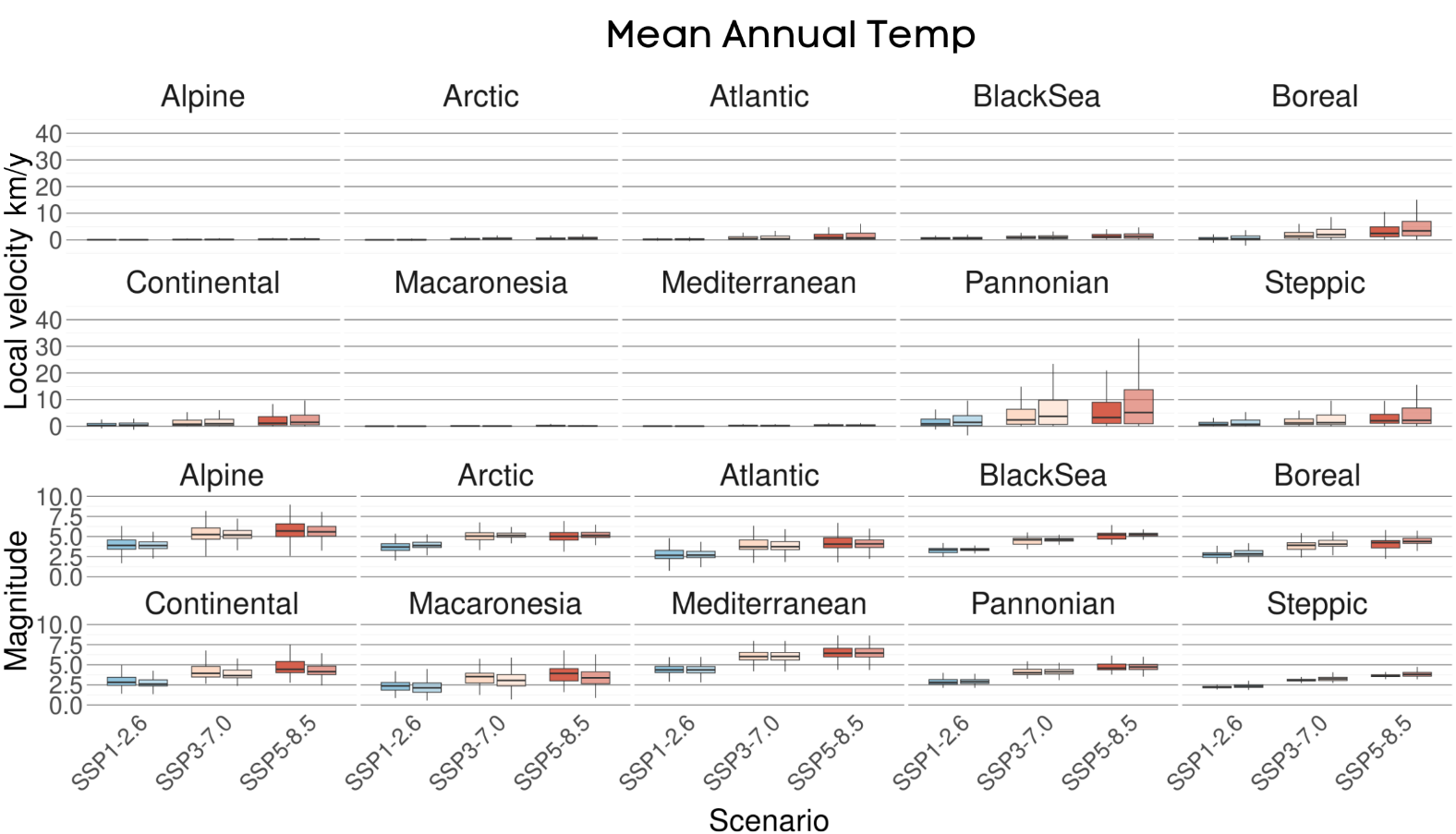

### Fig. S10) Boxplots oftemperature local velocity and magnitude in unprotected areas (darker colour) and PAs (lighter colour). Separate plots are shown for the 10 biomes covering the whole study area.

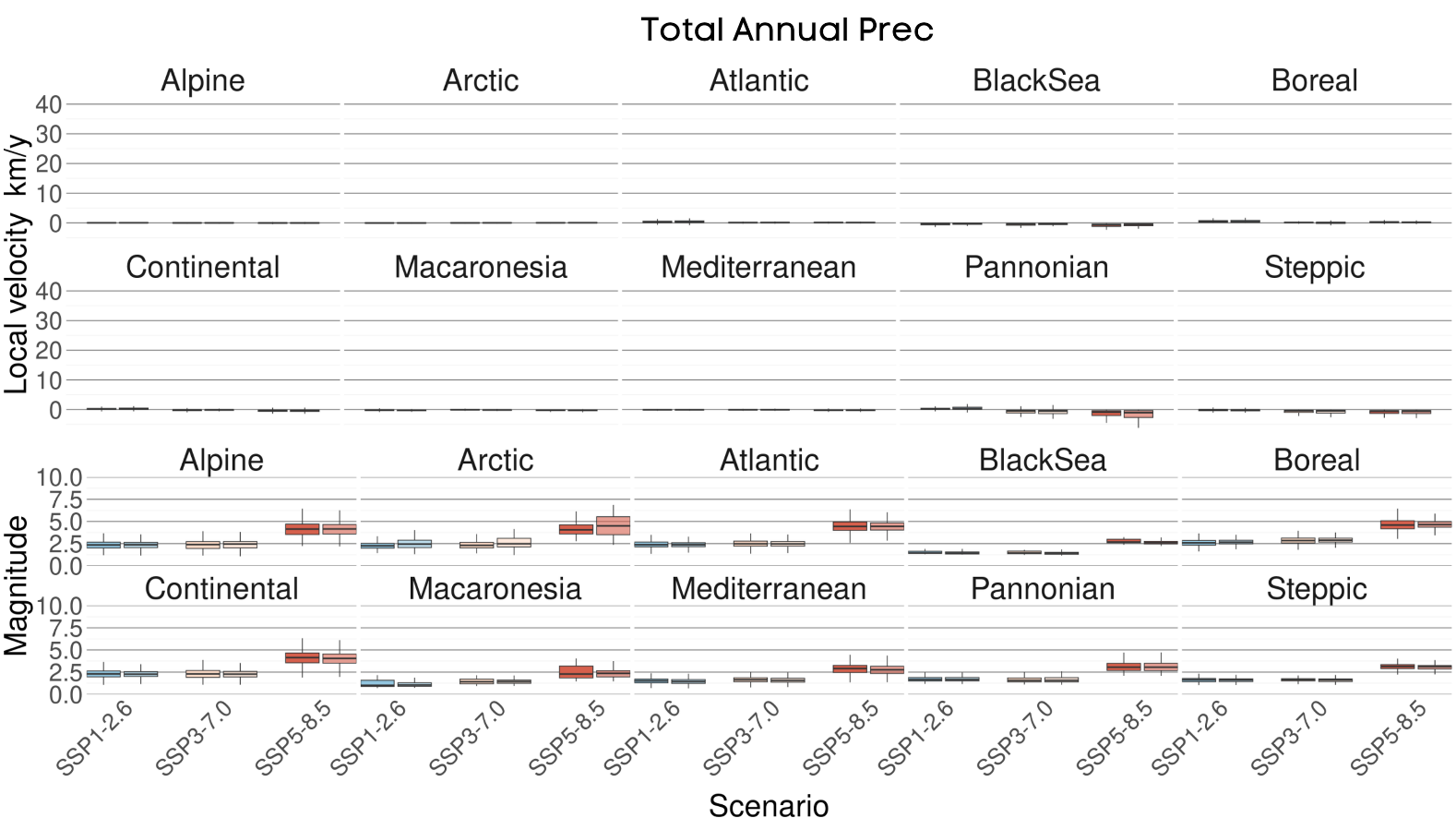

### Fig. S11) Boxplots ofprecipitation local velocity and magnitude in unprotected areas (darker colour) and PAs (lighter colour). Separate plots are shown for the 10 biomes covering the whole study area.

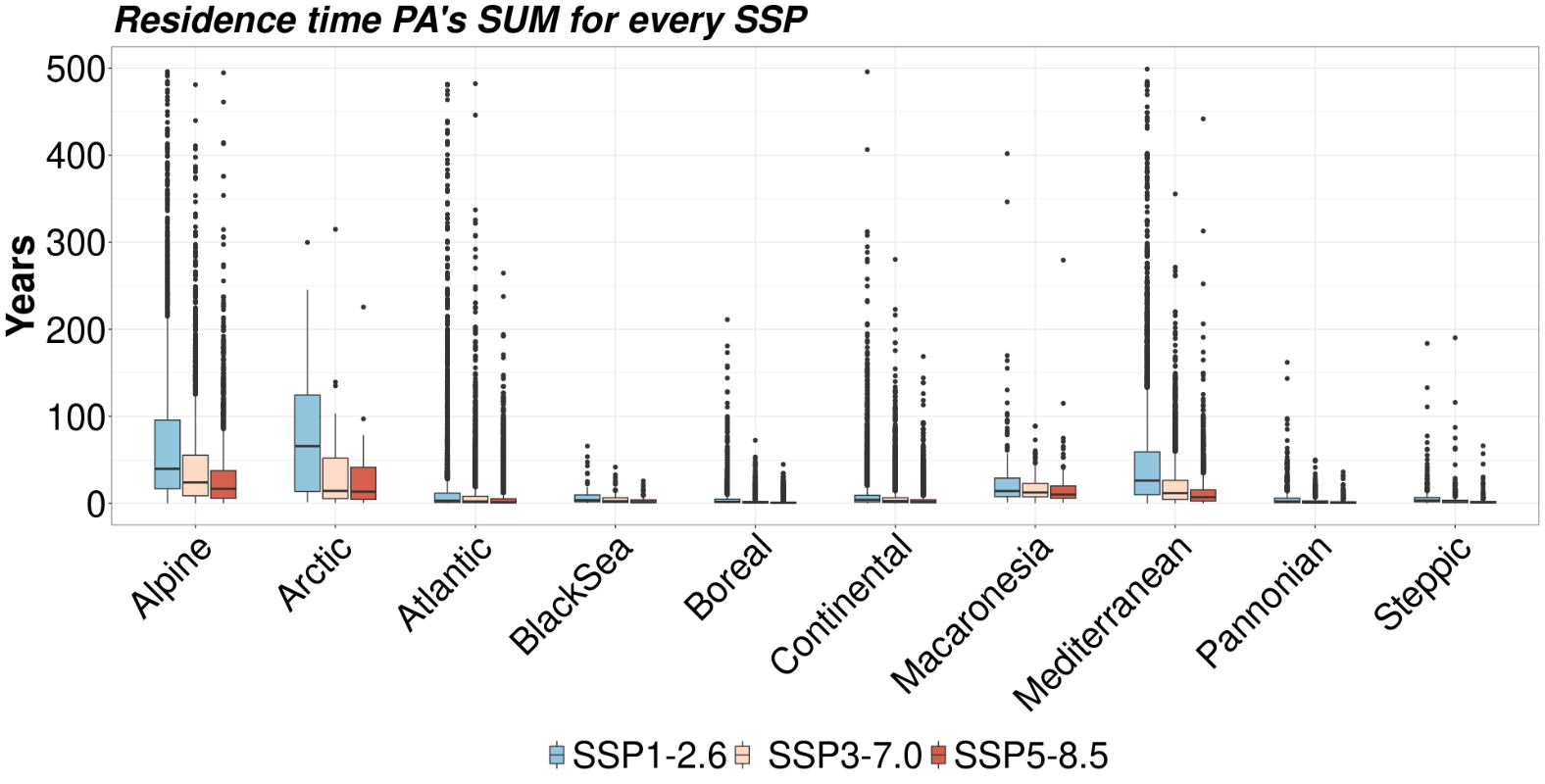

### Fig. S12) Projected residence time of European PAs based on climate local velocity for each biome, based on each analyzed SSP. Part of the outliers were cut for visualization purpose.

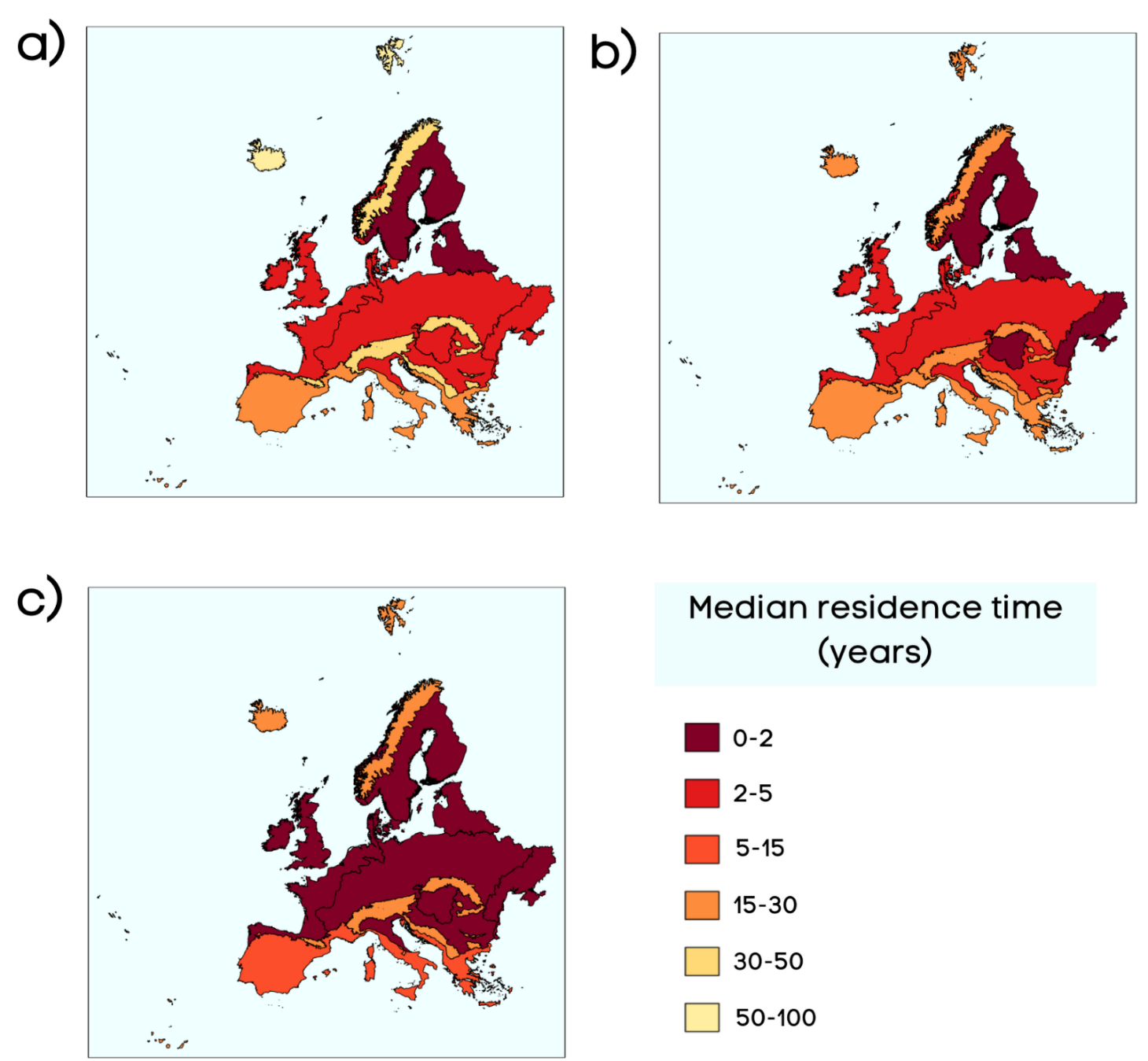

### Fig S13) Median climate residence time for PAs’ related to biomes under scenarios a) SSP1-2.6, b) SSP3-7.0, and c) SSP5-8.5.

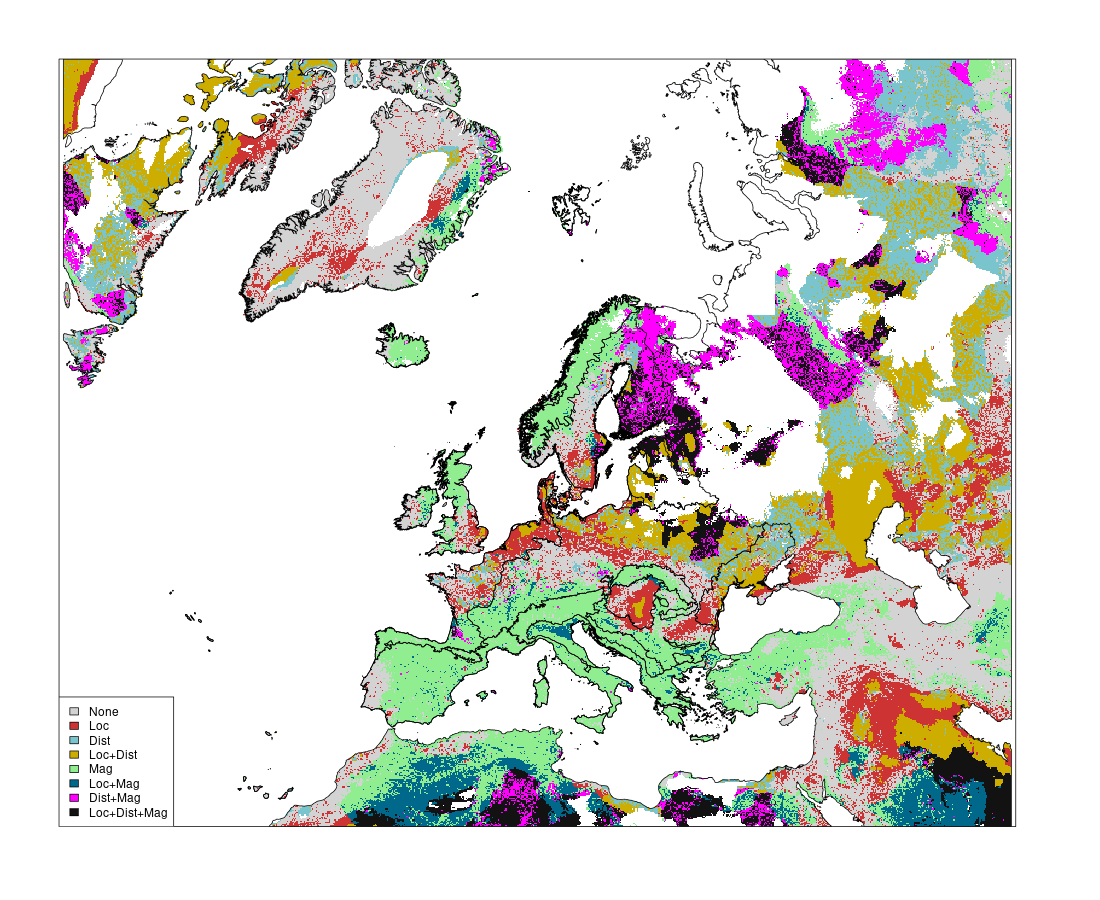

### Fig. S14) Overlay of the top-30% values from each climate risk metric for scenario SSP1-2.6. Areas in the map are color-coded based on their overlap with top-30% hotspot of exposure identified basedon different climate metrics (Loc, local velocity; Dist, distance velocity; Mag, magnitude).

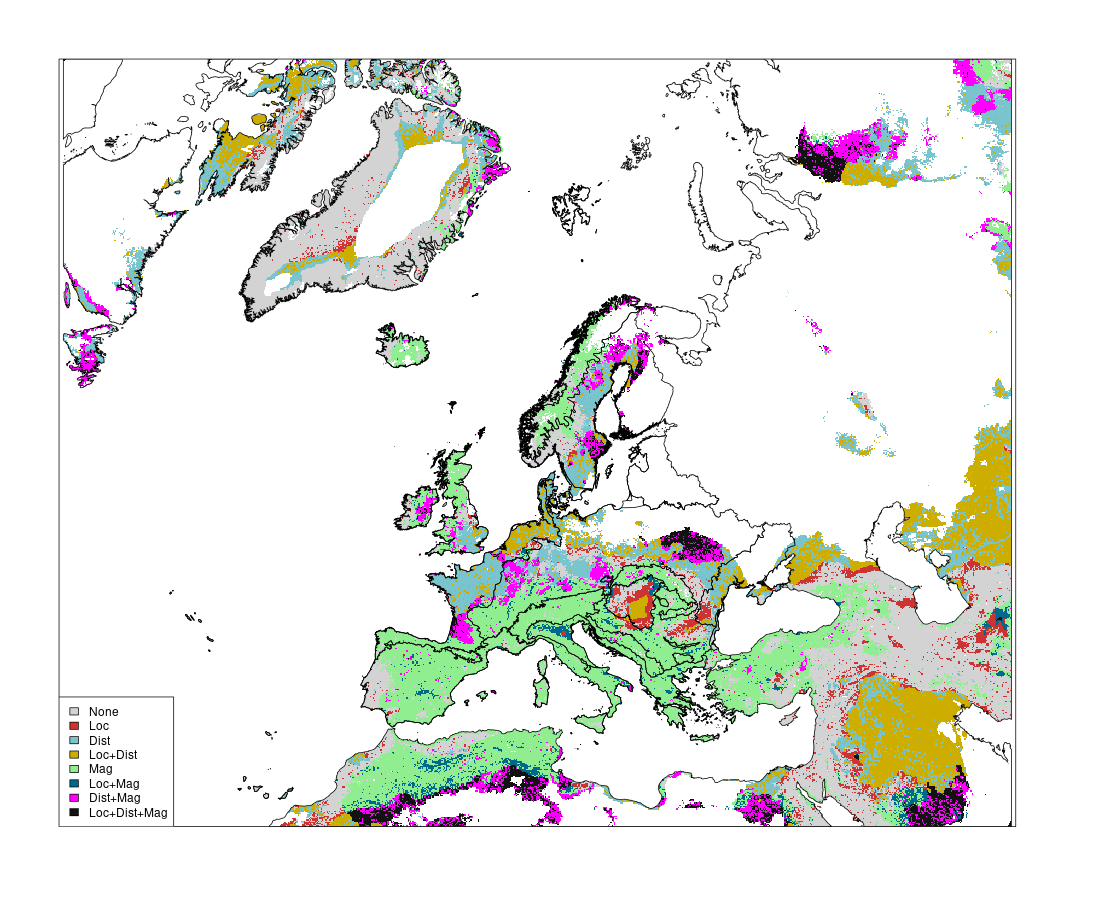

### Fig. S15) Overlay of the top-30% values from each climate risk metric for scenario SSP5-8.5. Areas in the map are color-coded based on their overlap with top-30% hotspot of exposure identified based on different climate metrics (Loc, local velocity; Dist, distance velocity; Mag, magnitude).

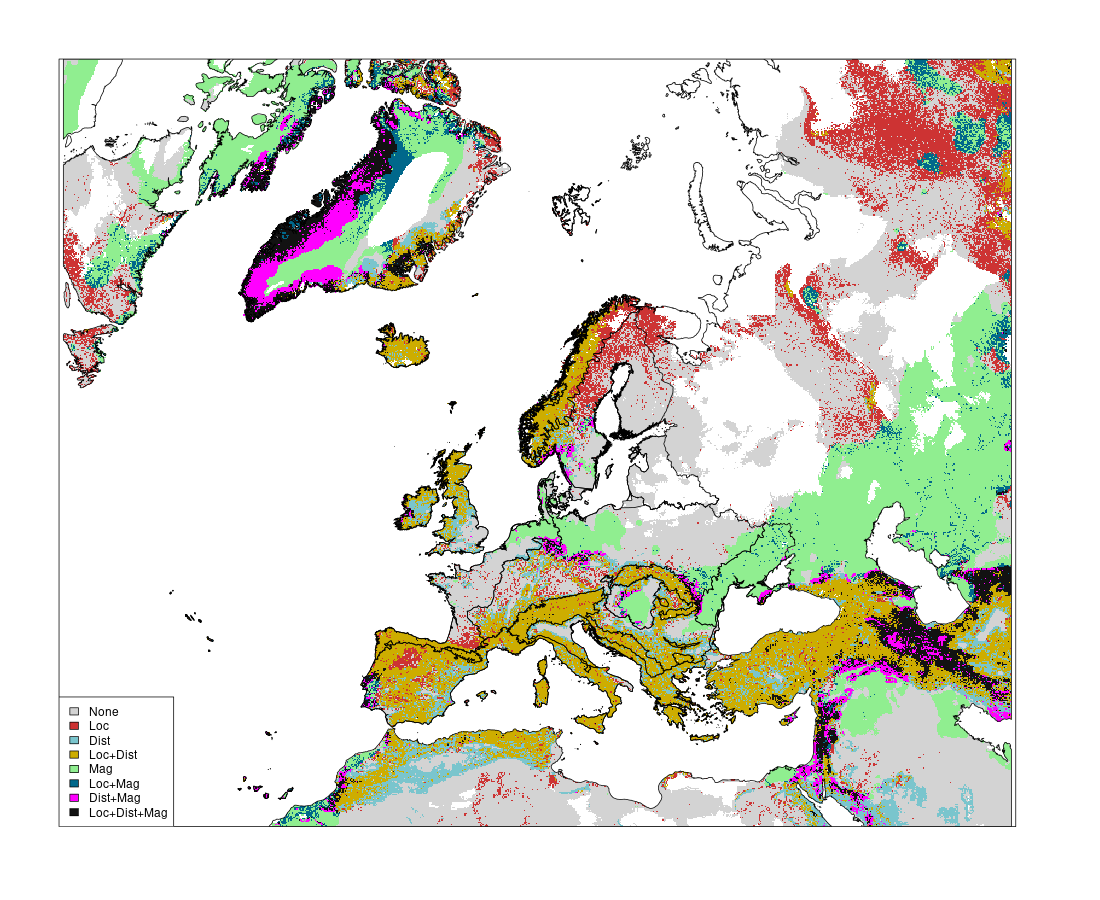

### Fig. S16) Overlay of the top-70% values (cold spots) from each climate risk metric for scenario SSP1-2.6. Areas in the map are color-coded based on their overlap with top-30% hotspot of exposure identified based on different climate metrics (Loc, local velocity; Dist, distance velocity; Mag, magnitude).

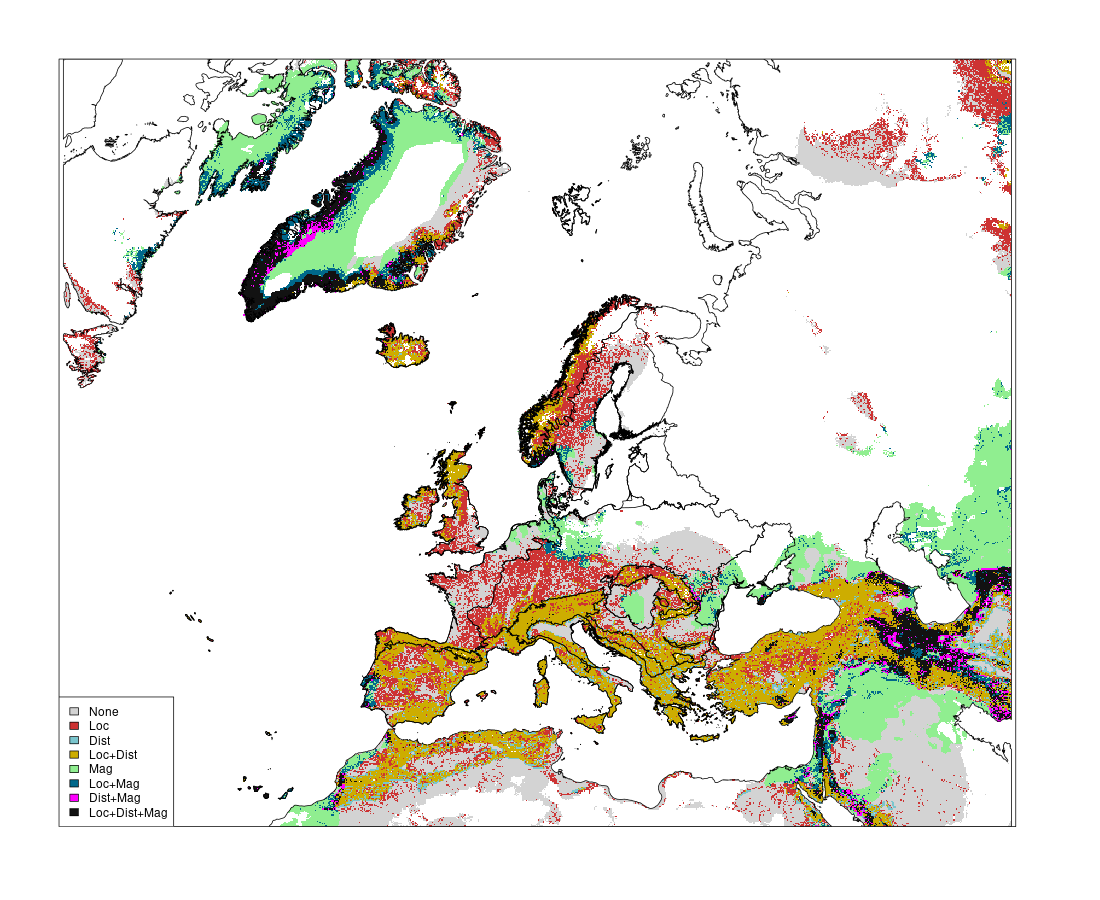

### Fig. S17) Overlay of the top-70% values (cold spots) from each climate risk metric for scenario SSP5-8.5. Areas in the map are color-coded based on their overlap with top-30% hotspot of exposure identified based on different climate metrics (Loc, local velocity; Dist, distance velocity; Mag, magnitude).

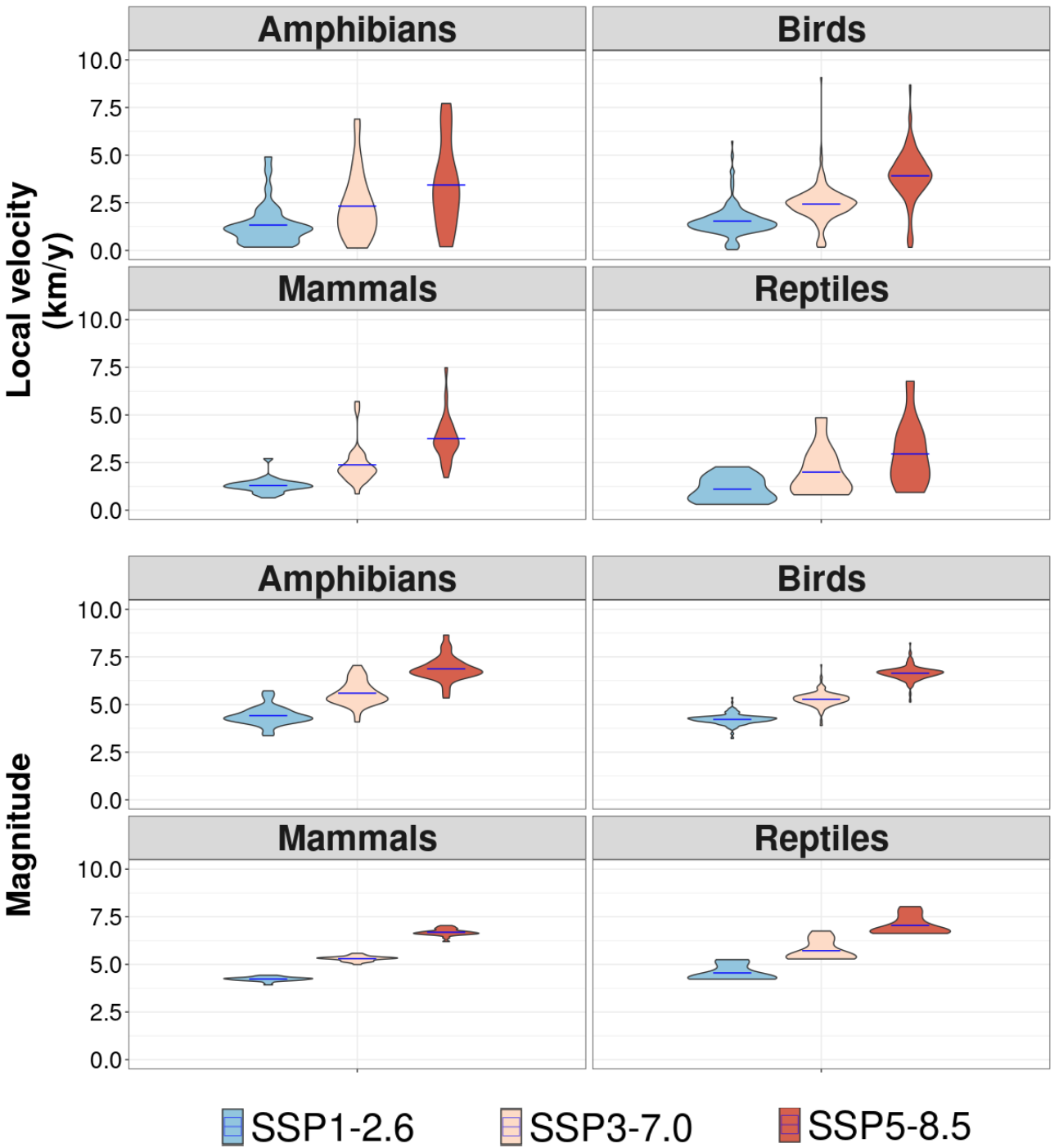

### Fig. S18) Predicted local velocity, distance velocity, and magnitude for the 1011 species occurring within Natura 2000 areas.

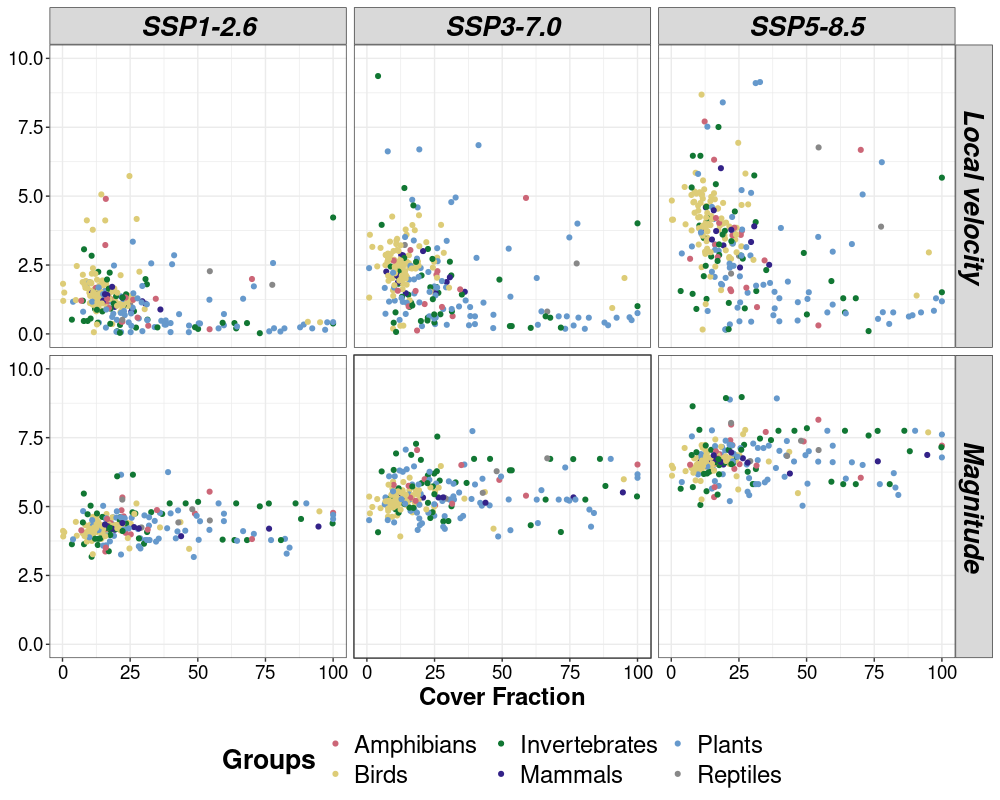

### Fig. S19) Predicted historical, SSP1-2.6, and SSP 5-8.5 climate metrics compared to the cover fraction of all species’ ranges located inside Natura 2000 areas (514 species).
